# Normalizing and denoising protein expression data from droplet-based single cell profiling

**DOI:** 10.1101/2020.02.24.963603

**Authors:** Matthew P. Mulè, Andrew J. Martins, John S. Tsang

**Affiliations:** Multiscale Systems Biology Section, Laboratory of Immune System Biology, National Institute of Allergy and Infectious Diseases (NIAID), National Institutes of Health (NIH); NIH Center for Human Immunology (CHI), National Institutes of Health (NIH); NIH-Oxford-Cambridge Scholars Program, Department of Medicine, Cambridge University

**Keywords:** Single cell, CITE-seq, normalization, gene expression

## Abstract

Multimodal single-cell protein and transcriptomic profiling (e.g. CITE-seq) holds promise for comprehensive dissection of cellular heterogeneity, yet protein counts measured by oligo-conjugated-antibody can have substantial noise that masks biological variations. Here we integrated experiments and computational analysis to reveal two major noise sources: protein-specific noise from unbound antibodies and cell-specific noise captured by the shared variance of isotype controls and background protein counts. We provide an open source R package (dsb) to denoise and normalize CITE-seq data based on these findings. (https://cran.r-project.org/web/packages/dsb/index.html).

## Introduction

A recent development in multi-omic single cell analysis involved using DNA barcoded antibodies to simultaneously profile surface proteins and transcriptome in single cells^1,2^. This greatly enhances our ability to discover, define, and interpret cell types and states, particularly those comprising the immune system given extensive existing knowledge connecting surface protein profiles to immune cell subsets and functions. Sequencing based assessment of surface protein expression bypasses spectral interference inherent in fluorescence-based cytometry methods, thus enabling simultaneous profiling of hundreds of proteins in single cells. Raw surface protein count data from such approaches, for example, cellular indexing of transcriptomes and epitopes by sequencing (CITE-seq), are typically positive and non-sparse (<1% to 14% zero in datasets analyzed here), in contrast to droplet-based single cell mRNA data, (typically >90% zero), resulting in low-level denoising and cell-to-cell normalization challenges distinct from both continuous cytometry protein expression and single cell mRNA data. While low-level normalization and modeling approaches for single cell RNA-seq data have received considerable attention^3–8^, those for protein count data are in their infancy despite the need for such approaches given the substantial levels of noise reported in raw protein counts^2^. Here we performed experiments and computational analyses to reveal two major components of protein expression noise in droplet-based single cell experiments: 1) protein-specific noise originating from ambient, unbound antibody encapsulated in droplets that can be accurately inferred via the average protein levels detected in empty droplets, and 2) droplet/cell specific noise revealed via the shared variance component associated with isotype antibody controls and background protein counts in each cell. We developed a software package, “dsb”, the first low-level normalization method developed specifically for this type of protein count data, to correct for these noise sources. Application of this approach to both our own and multiple external data sets demonstrated its generalizability to enhance downstream analysis, including manual and automatic identification of cell populations and states.

### Analysis of unstained cells reveals ambient antibody capture as a major source of protein-specific noise

To assess protein count noise, we utilized our previously reported dataset measuring more than 50,000 Peripheral Mononuclear Cells (PBMCs) from 20 healthy human donors^9^ stained with an 87 CITE-seq antibody panel (including four isotype controls; Totalseq-A reagents, Biolegend). Consistent with the original CITE-seq report^2^, we noticed non-zero counts for most proteins in each cell, resulting in positive counts even of markers not expected to be expressed in certain cell types. We also noticed non-zero, “ambient” protein counts in tens of thousands of empty droplets containing capture beads without cells, which emerge naturally due to Poisson distributed cell loading, reminiscent of cell-free RNA observed in droplet-based single cell RNAseq^10^. We reasoned that background noise in CITE-seq data may partly reflect such unbound, ambient antibodies captured in droplets. To assess whether counts in empty droplets indeed reflect the ambient component in cell-containing droplets, we compared background protein levels in cell-free droplets with droplets capturing unstained control cells spiked into the cell mixture after cell staining and washing but prior to droplet generation (Fig. 1a). We found positive protein counts even for unstained control cells, and that the average log transformed level per protein in empty droplets and unstained control cells were highly correlated (Fig. 1b). A similarly strong correlation was observed between the average protein counts in stained cells “negative” for the protein and those in empty droplets (Supplementary Fig. 1a-c; “negative” cells correspond to those in the fraction with lower expression of the protein; two fractions of cells were inferred by Gaussian mixture models - see Methods), further suggesting that the noise component correlated across cells is dominated by ambient antibody capture. Thus, protein counts in empty droplets, which are available in all single cell droplet experiments, provide a direct estimate of the ambient background due to free antibody capture for each protein. This observation motivated the first step of our dsb normalization method to transform counts of each protein in cell-containing droplets by subtracting the mean and dividing by the standard deviation of that same protein across empty droplets (see Methods). The resulting transformed protein expression index in cell-containing droplets reflects the number of standard deviations above the expected ambient capture noise, thus centering the negative cell population for each protein around zero to help improve interpretability of the resulting protein expression values.

**Figure 1.**
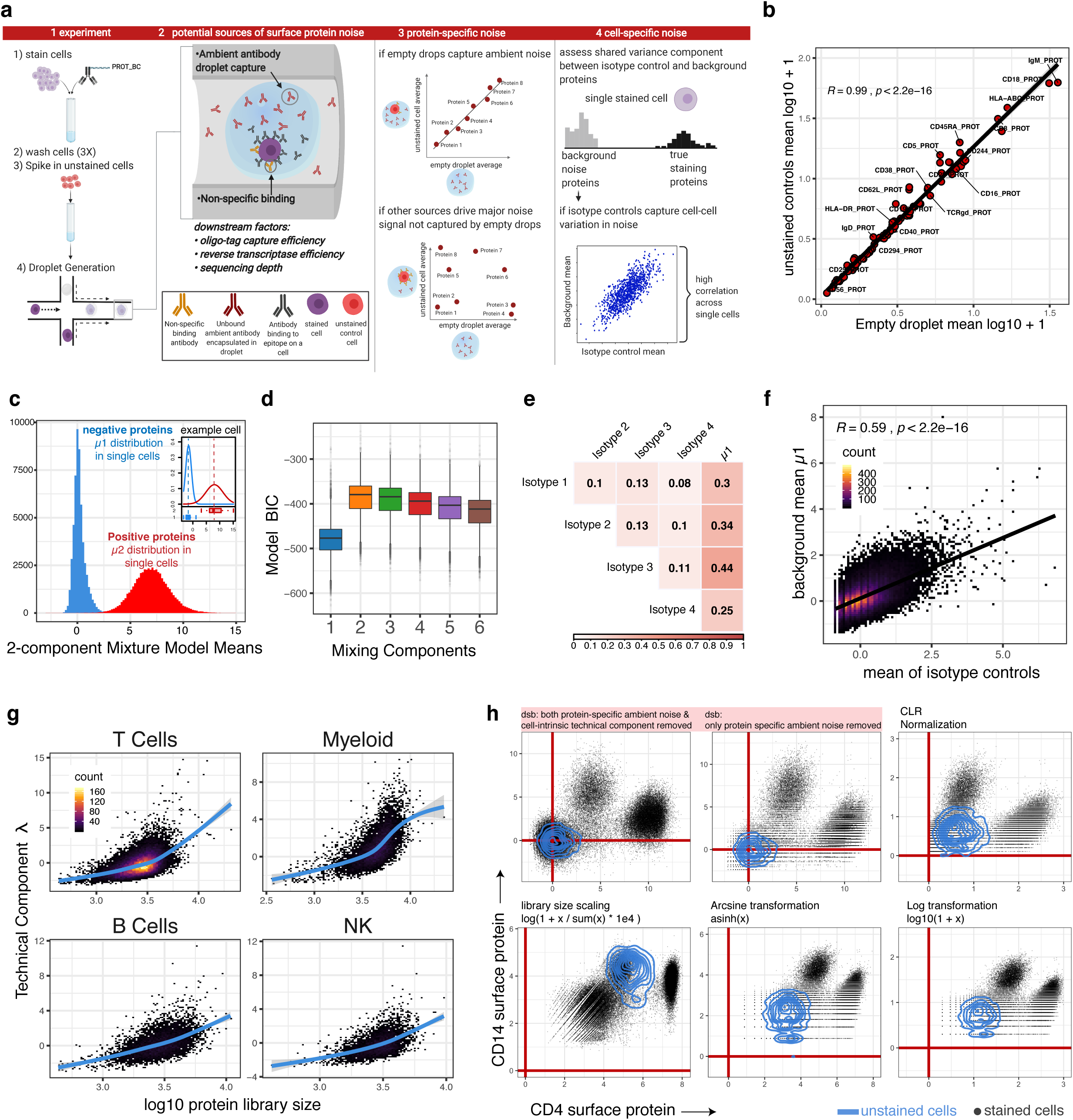
Protein noise source assessment. **a.** 1-2: Schematic illustrating experimental setup and potential sources of noise in CITE-seq data; 3: assessing protein-specific noise: expected outcomes from unstained cell experiment to evaluate whether unbound, ambient antibody encapsulated in droplets constitute a major source of protein-specific noise in CITE-seq data. If empty droplets capture the ambient component of background noise their counts should be highly correlated with unstained control cells (top). If control cells contain information on noise sources not captured by empty drops, the correlation should be weak. 4: assessing cell-specific noise by evaluating the correlation between background counts with isotype control signals across single cells. **b.** Scatter plot of the average protein log10(count+1) of unstained control cells spiked into the stained cell pool prior to droplet generation (y-axis) versus that of empty droplets (x-axis) without a cell. Pearson correlation coefficient and p value are shown. Cells negative for each protein have a similar correlation structure with empty droplets (Supplementary Figs. 1a-b). **c.** A two-component Gaussian mixture model (i.e., with k=2 subpopulations) was fitted to the protein counts in single cells (inclusive of all proteins/antibodies except isotype controls); shown are the distributions of the component means from all single cell fits (blue=“negative” population; red=“positive” population); data from an arbitrary randomly-selected cell is shown in the inset. **d.** Comparison of Gaussian mixture models fit to single cells with between k = 1 and k = 6 subpopulations (defined as “G” in the *mclust* package) vs the Bayesian Information Criteria (BIC) of the model fit; a total of n=6 models were fitted to the dsb normalized counts of each cell from step I (rescaling using background from empty droplets) but prior to step II denoising (see methods). Models were fit with the expectation maximization algorithm within the mclust package which defines the best fit as the model that maximizes the BIC (see methods and supplemental note); to reduce memory requirement for this analysis, the n = 28,229 cells from batch 1 were analyzed and the BIC and parameters from the resulting 169,374 models are shown in the figure. **e.** Pearson correlation coefficients among isotype controls and background component mean inferred by Gaussian mixture model (μ1, which was fitted per cell as in c); all corresponding p values are less than 0.001. **f.** Scatter density plot between μ1, the mean of each cell’s negative subpopulation from the per-cell Gaussian mixture model (blue in Fig. 1c) versus the mean of the four isotype controls across single cells. Pearson correlation coefficient and p value is shown. **g.** The first principal component of μ1 and isotype control vectors (five vectors in total) is used to define each cell’s “technical component” λ; the value of λ (with sign inverted to be consistent in direction) is plotted against the cell’s protein library size (number of UMIs) within each major cell lineage (see also Supplementary Fig. 3a for additional cell clusters). **h**. Single cell protein expression of CD4 vs. CD14 normalized by different methods. Contour lines in blue are the distribution of unstained control cell CD4 and CD14 normalized in the exact same way as the stained cells in black within each plot, including dsb normalization using the same empty droplets for ambient correction of the unstained cells. The first two panels from the top left correspond to 1) dsb normalization with both ambient (empty droplet) background subtraction and regressing out technical component λ; 2) same as previous but without regressing out the technical component; other transformations shown to the right have been used for single cell data (e.g., flow cytometry, CyTOF, scRNAseq), including the CLR transformation reported in the original CITE-seq publication are shown. Outlier cells (less than 0.3% of total cells in any panel) are removed to focus on the three main cell populations.

### Shared variances between isotype controls and background protein counts in single cells provide cell-intrinsic normalization factors

In addition to ambient noise correlated across single cells as captured by average readouts from empty droplets, cell-intrinsic technical factors, including but not limited to oligo tag capture, cell lysis, reverse transcriptase efficiency, sequencing depth and non-specific antibody binding, can contribute to cell-to-cell variations in protein counts that should ideally be normalized across single cells. Common normalization procedures for mRNA standardize the total number of reads between cells by calculating scaling factors based on each cell’s total “library size”^11^ (defined herein as the total UMI count in each cell). However, these methods are not appropriate for surface protein count data because: 1) current methods/experiments still measure only a small fraction of unique proteins with a wide range of antigen density on target cell types, resulting in individual protein counts in single cells of less than 10 to more than 1000; differences in total protein counts between cells therefore depend on the specific antibody panel used and may be dominated by differences between cell types. 2) Protein library size scaling factors could reflect biologically relevant variations, e.g. those due to the physical size of naïve vs. activated lymphocytes. Here we integrated two types of independently-derived measures to reveal a conservative (i.e., avoiding over-correction and removal of biological information), robust estimate of the factor associated with cell-intrinsic technical noise (Fig. 1a).

First, the four isotype control antibodies with non-human antigen specificities in our panel could in principle capture contributions from non-specific binding and other technical factors discussed above. The counts of the isotype controls were only weakly (but significantly) correlated with each other across cells (Fig. 1e), suggesting that while each may be individually noisy, their shared component of variation could capture technical noise in the experiment. Indeed, the correlation between the mean of four isotype controls and the protein library size across single cells was higher (Pearson correlation 0.45) than that of individual isotype controls (average Pearson correlation 0.22). Second, to further boost the robustness of estimating cell-intrinsic technical noise, particularly given that the number of isotype controls available in practice can be limited, we sought an additional method to estimate this technical variation. We reasoned that since each cell in a sample of multiple distinct cell types (e.g. PBMCs) is expected to express only a subset of protein markers in staining panels, each cell should exhibit a bimodal protein count distribution comprising “negative” and “positive” populations. If so, the average counts in the negative population of non-staining proteins, e.g., those specifically expressed in another cell lineage, could reflect and therefore serve as another readout of the cell’s technical component and can then be integrated with noise measured by isotype controls. To assess this bimodal distribution assumption, we applied a Gaussian mixture model with two (k=2) subpopulations to fit the protein counts within each single cell after correcting for the protein-specific ambient noise we identified above (see below and Methods; Fig. 1c). We found clear separation between the background (with mean=μ1) and positive (mean=μ2) population with substantial cell-to-cell heterogeneity of subpopulation means (Fig. 1c). Two-component models had the best fit in a majority (81%) of cells when comparing k=1 to 6 component models assessed using the Bayesian Information Criterion (BIC) ([see Supplementary Note] Fig. 1d, Supplementary Figs. 2a-b). While k=3 models had the best fit in nearly all remaining cells, they had relatively similar BIC as the corresponding k=2 models (Supplementary Fig. 2c), indicating that the 2-component fits were identifying similar positive and negative populations (Supplementary Fig. 2d–f). These data indicate that a single, negative background protein count population can be clearly identified for most cells. The bimodal distribution assumption also held well in our assessment of five different datasets with a range of proteins profiled (from greater than 80 to just 14, Fig. 1d and see below).

We next assessed information sharing between μ1 and the isotype control mean. The correlations between μ1 and each individual isotype control (average correlation r = 0.33) or the average of all four isotype controls (r=0.59; Fig. 1f) were higher than those between the isotype control themselves (average correlation r = 0.11), suggesting that the shared variation (i.e., average) between the independently inferred μ1 and isotype controls captured unobserved, latent factors contributing to technical noise (Figs. 1e-f). We thus reasoned that the first principal component (λ) capturing the shared variation of μ1 and the isotype controls would be a robust measure of cell intrinsic technical noise. Supporting this idea, λ was associated with the protein library size *within* each cell lineage and protein defined cell type (Fig. 1g and Supplementary Fig 3a clusters defined after dsb steps I and II, see Methods). Given this information sharing between μ1 and isotype controls, we recommend inclusion of multiple isotype controls in CITE-seq experiments to serve as anchors for revealing technical normalization factors (see Supplementary Note). Together, our data indicate that while the signal from individual measures such as isotype controls can be noisy and may reflect multiple yet often unknown sources of variation, their correlated component of variation can serve as a robust normalization factor for surface protein expression in single cells. Thus, in a second, optional step dsb regresses out λ from single cells after the ambient noise correction in step 1 (see Methods).

### Application of dsb in different datasets

We next applied dsb to both our own and external, independent data sets to assess its utility in practice. Overall, dsb normalized values improve identification of discrete cell populations positive for markers above zero-centered background noise (Fig. 1h). dsb normalized protein values can facilitate robust manual gating across immune cell lineages (Supplementary Fig. 4a), and unbiased identification of cell populations via protein-based clustering. Distinguishing non-zero marker expression from noise within individual cell clusters can be challenging when using transformations such as the centered log ratio (CLR) because, for example, all CLR normalized protein values for cells in cluster 4 (framed cluster in Fig. 2a) are non-zero and therefore cells appear to express markers known to be specific for other cell lineages, such as IgA/IgM (B cells), CD16 (Monocytes / NK cells) and CD57 (T and NK cells) (Fig. 2b). In contrast, dsb normalized values for IgA, IgM, and CD57 are zero-centered (Fig. 2b), as the level of these proteins in this cluster is statistically similar to the level in empty droplets (Figs. 2c and 2d–red proteins) while CD16, CD244, and CD56 have higher dsb values (greater than 8 standard deviations above the mean in empty droplets, +/- the correction from regressing out the technical component). This cluster is thus consistent with the known phenotype of CD57 negative CD16++CD56+ NK cells, which are not known to express B-cell specific markers such as IgM or IgA. Clusters identified based on dsb normalized protein values had biologically coherent expression of cell type-defining proteins (Fig. 2e, f, Supplementary Figs. 4b–c, See ref. 9).

**Figure 2.**
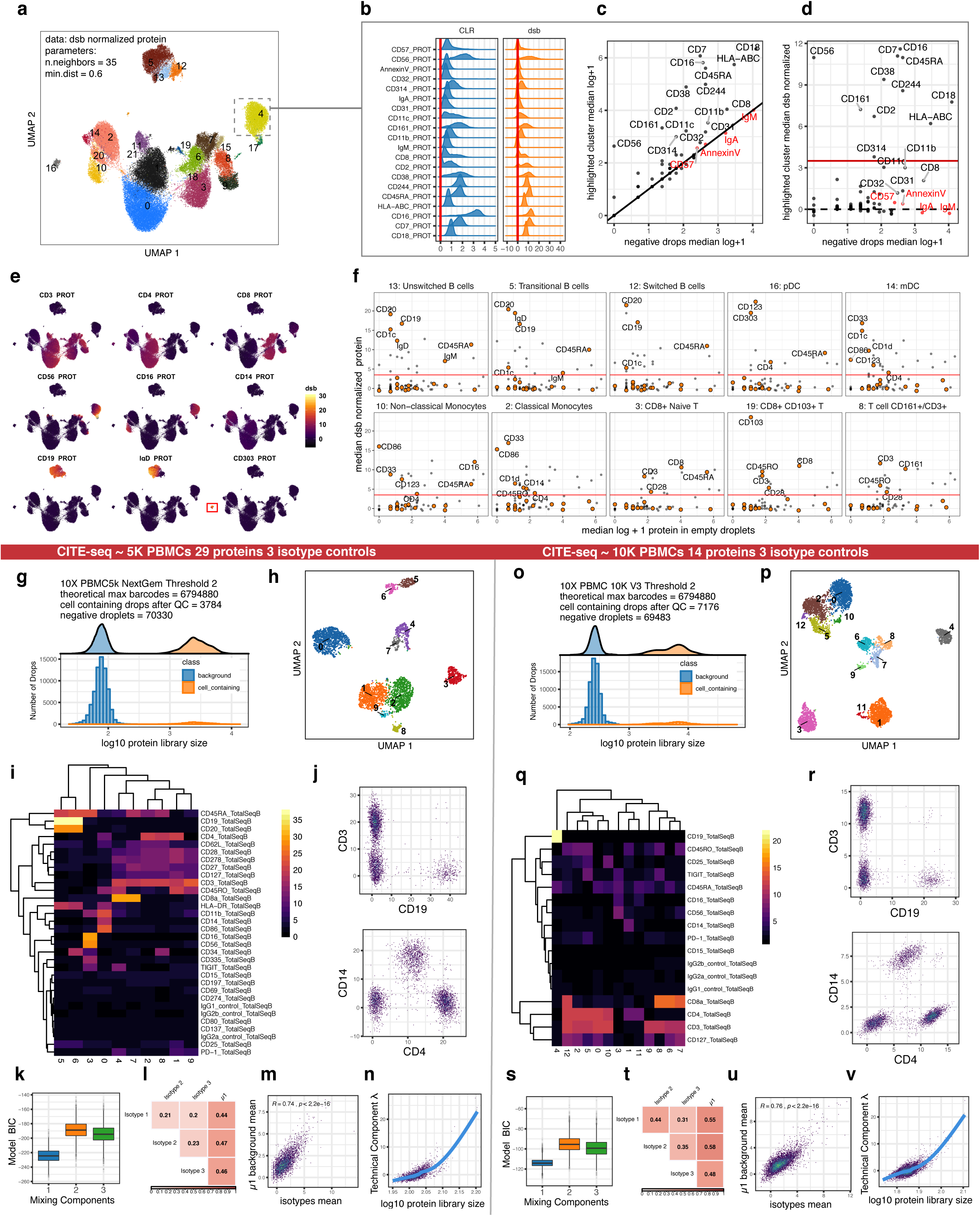
Application of dsb normalization for manual gating, automatic population identification, and in external datasets. **a**. UMAP embeddings of single cells labeled by cluster derived from clustering on dsb normalized protein levels. **b**. The distribution of protein expression of cluster 4 (highlighted with a grey box in (a)) using CLR or dsb for normalization. **c**. Median log + 1 protein levels in cells from cluster 4 versus empty droplets; proteins highlighted in red are comparable in expression to “positive” proteins after log transformation but are similar to background levels in empty droplets. All proteins with median log expression greater than 2.4 are labeled with the protein name. **d.** Similar to (c), but the y axis shows the median dsb normalized values; proteins in red (those near the diagonal in (c)) are now residing around zero reflecting their proximity with mean counts in the empty droplets; proteins above the red line have median dsb normalized expression within the highlighted cluster 4 (see (a) and (b)) above 3.5 i.e., 3.5 standard deviations above ambient noise, +/- adjustment for the cell intrinsic technical component. **e.** UMAP embeddings as shown in (a) with overlay of dsb normalized expression for a subset of lineage-defining proteins–each protein is visualized on the same continuous dsb normalized units corresponding to the number of standard deviations from the mean of the empty droplets **f.** as in d, with a subset of informative proteins for cluster identification highlighted in orange in each panel for comparison of the behavior of marker expression between clusters; those with a dsb value above 3.5 (red line) are annotated with the protein name within each panel. The proteins highlighted in orange in each panel are CD20, CD19, IgD, IgM, CD123, CD303, CD1d, CD1c, CD14, CD103, CD16, CD3, CD4, CD8, CD28, CD161, CD45RO, CD45RA, CD33, and CD86. Panels **g-v**, data downloaded from support.10xgenomics.com for assessing the applicability of dsb normalization to independent datasets. Panels **g-n**: applying dsb to a publicly available dataset generated using 10X genomics “NextGem” chemistry measuring 29 proteins across ~5K cells. **g.** Distribution of protein library size of a subset (see Supplementary Note and Supplementary Fig. 8) of the raw cell ranger output showing bimodal peaks: 1) a background peak consisting of ~50,000 barcodes and a second peak of ~3700 cells after mRNA based QC (see Methods and software tutorial). **h.** UMAP of cells based on dsb normalized protein counts; cells were clustered using surface proteins only and are colored by cluster identity. **i.** Heatmap of the average expression of dsb normalized protein (rows) vs. clusters (columns). **j.** Biaxial plots of single cells using canonical T cell, B cell, and monocyte markers: T cells (CD3+ CD19-), B cells (CD3-CD19+), monocytes (CD4 mid CD14 high), and CD4 T cells (CD14-CD4+) are clearly distinguishable. **k.** As in Supplementary Figure 2, the distribution of BICs of Gaussian mixture models fitted to single cells with 1-3 components. **l**. Similar to Fig. 1e showing the correlation among isotype controls and mean of background protein counts across single cells. **m.** As in Fig. 1f, the correlation between the mean of isotype controls and the mean of the background population across single cells. **n.** Similar to panels in Fig. 1g showing the relationship between the inferred technical component and protein library size across single cells. Panels **o-v**: Identical kinds of panels as g-n but using data from 10X Genomics V3 chemistry with 14 proteins across ~10K cells. See also dsb normalization assessment using additional independent datasets in Supplementary Fig. 5: data from 10X Genomics “5 Prime” chemistry, Supplementary Fig. 6: data from 10X Genomics Chromium Version 3 “PBMC 5K data”, and Supplementary Fig. 7: data from the Mission Bio Tapestri platform for combined surface protein and genotype analysis.

Using several independent, publicly available CITE-seq datasets, we next tested the general applicability of dsb and assessed whether the modeling assumptions developed using our own data would generalize to these datasets. External datasets were generated using versions of the 10X Genomics droplet profiling kit distinct from the one we used^9^ and they profiled 14–29 surface proteins in ~5,000 to 10,000 cells. Similar to our dataset, we detected a large number of empty droplets (>50,000) from which to estimate the protein-specific ambient noise (cell-containing barcodes and background/empty droplets were clearly evident as the upper and lower peaks in protein library size distribution, respectively - see discussion in Supplementary Note); the number of cell-containing droplets recovered after quality control (3,000-8,000 droplets) was also consistent with the number of loaded cells (Figs. 2g, 2o, Supplementary Figs. 5a, 6a).

Applying dsb without further modification resulted in biologically interpretable protein-based clusters (Figs. 2h,i,p,q, Supplementary Figs. 5b-d 6b-d); canonical immune cell populations could be clearly delineated by conventional biaxial plots (Figs. 2j, 2r, Supplementary Figs. 5e-g, 6e-g). Importantly, the model fitting behavior and correlations among isotype controls and background counts observed in our dataset were similarly observed in these independent datasets: 1) The k=2 component Gaussian mixture model had the best fit according to BIC in most single cells (89% average across n=4 datasets; see Figs. 2k, 2s, Supplementary Figs 5h, 6h); 2) the estimated μ1 (mean of background protein counts) for each cell corelated significantly with the mean of isotype controls across single cells (Figs. 2l-m, 2t-u, Supplementary Figs. 5i-j, 6i-j); finally, 3) the inferred technical component using isotype controls and μ1 was correlated with the protein library size. (Figs. 2n, 2v, Supplementary Figs. 5k, 6k). In addition to the 10x platform, we further tested applying the dsb package to simultaneous single cell surface protein and DNA measurements from ~1,000 cells (MissionBio Tapestri) (Supplementary Fig. 7a-e) and found that our normalization is similarly applicable and can help improve interpretation of cell cluster identities after correcting for ambient background estimated from >16,000 background droplets.

## Discussion

The developers of CITE-seq accounted for protein-specific noise in human cord blood cells^2^ by spiking in mouse cells to set a human-specific cutoff for determining whether a CLR transformed protein expression value is higher above noise. This approach entails more complex experiments and downstream analyses; some subsequent reports of CITE-seq have used CLR transformation without control cells^12,13^. The dsb approach mitigates the need for control cells from another species. Recent methods proposed to use joint probabilistic models of mRNA and protein^14–16^ to model logarithmic or CLR transformed protein values, with one of the goals being the identification of protein expression above noise. For example, TotalVI^15^ uses an mRNA and protein generative neural network model to estimate posterior probability distributions of protein expression, which identified cells with zero, low, or high probability of CD4 protein level in human PBMCs. As expected, this identified monocytes and T-helper cells based on known low and high surface CD4 protein levels on these cells, respectively; these populations were similarly recovered by dsb normalized populations centered at 0, 4, and 10 (Fig 1h and see Fig 2j, r Supplementary Fig. 5f, 6f). Such joint machine learning models are complementary to and distinct from dsb, which focuses on low-level denoising and normalization challenges unique to CITE-seq protein data by directly inferring and removing the two noise components delineated in our experimental and computational analyses above. The resulting denoised and normalized data from dsb can be used in any downstream analysis; dsb is thus compatible with approaches that focus on higher level single cell data analysis, such as joint protein-mRNA models and clustering methods^17^. The dsb package is computationally efficient and can process ~10^5^ cells on a laptop, e.g., the primary dataset in this study (with >53,000 cells) was normalized and denoised in under 4 minutes. The output can be easily integrated with diverse single cell software platforms such as Bioconductor^18^, Seurat^19^, and scanpy^20^.

## Acknowledgements

This work was funded by the intramural research program of NIAID, NIH. The authors thank Can Liu for testing the dsb package and Yuri Kotliarov for helpful discussions and data deposition assistance related to this work. Illustrations were created with Biorender (biorender.com).

## Author Contributions

MPM, AJM and JST designed experiments, devised analysis strategies, interpreted data, and developed the dsb method. AJM and MPM performed CITE-seq experiments. MPM created the dsb R package and performed analysis. All authors contributed to the writing of the manuscript.

## Methods

### The denoised scaled by background normalization (dsb) method

The dsb method is implemented through the R package “dsb” through the function “DSBNormalizeProtein” which models and accounts for 1) protein-specific ambient noise correlated across single cells as captured by average readouts from empty droplets and 2) droplet/cell specific technical noise revealed via the shared variance component associated with isotype control antibodies and background protein counts in each cell. In step I, protein counts in empty droplets are used to estimate the expected ambient background noise for each antibody. Each protein’s counts in cell-containing droplets are thus rescaled using this expected noise measurement as:

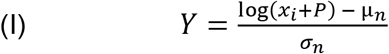

Where log *x_i_* is the natural log of the count for protein Y in cell *i, P* is a pseudocount added to prevent taking the log of zero and to stabilize the variance of small counts, and *μ_n_* and *σ_n_* are the mean and standard deviation of empty droplets for protein Y, respectively, computed in the same way in natural log space with pseudocount *P* added. The value of *P* can be empirically chosen; we use 10 by default, finding this provides good clustering performance and visualization of the CITE-seq data we have analyzed. The transformed expression estimate (Y) for the protein in each cell can be interpreted as the number of standard deviations above the expected ambient background noise of that protein.

In step II, dsb denoises cell-to-cell technical variations by defining and removing the “technical component” of each cell’s protein values after ambient correction from step 1. In this optional but recommended step, dsb first fits a Gaussian mixture model through the expectation-maximization algorithm implemented with the mclust(1) R package to the transformed count of each cell from step 1 with k = 2 mixture components:

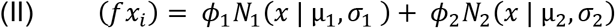

In the model above, each cell’s log normally distributed proteins comprising the nonstaining noise/background protein subpopulation for each cell are estimated by (N1). dsb then combines μ1, representing the mean of the background protein subpopulation N1 in that cell, and the cell’s isotype control antibodies by calculating principal component 1 (i.e. the primary latent component “λ”) through these variables; this defines the cell’s “technical component”. The technical component for each cell is then regressed out of the ambient-noise-corrected values from part 1 (i.e. dsb internally uses limma(2) to subtract the coefficient for λ of a linear model fitted to the ambient-corrected values). See Supplementary Note for additional information on usage and definition of the technical component in experiments without isotype controls.

### CITE-seq on 20 human PBMC samples

CITE-seq data analyzed here were previously used to assess the cellular origin and circuitry of baseline immune signatures(3); an earlier version of dsb was used therein to normalize the protein data which is identical to the default method implemented in the dsb package, with exception of the pseudocount used (1 vs 10, see below). Experiment details can be found in our prior report(3). Briefly, oligo-labelled antibodies for sample barcoding (cell “hashing”) and surface target protein detection were obtained from Biolegend. After incubating each sample with a barcoding antibody(4), cells from each donor were pooled into one tube and stained with an optimized mixture of oligo-labelled CITE-seq antibodies against target surface proteins. Two experimental batches were performed on consecutive days, using aliquots of the same pool of antibodies for each batch. The pooled donor cells from each of two batches were each distributed evenly across 6 lanes (per batch) of the 10x Genomics Chromium Controller using Single Cell 3’ expression reagents (version 2). Sample barcoding (HTO) and target surface protein (ADT) libraries were prepared as in the original CITE-seq report and according to the publicly available CITE-seq protocol (version 2018-02-12, cite-seq.com). cDNA libraries were prepared using the 10x Genomics v2 kit according to manufacturer’s instructions. Libraries were sequenced using the Illumina HiSeq 2500 using v4 reagents. We used CITE-seq Count(5) for HTO and ADT read mapping and Cell Ranger for RNA mapping, and cells were then demultiplexed as previously reported(3,4,6) (see Supplementary Note for additional details on demultiplexing).

### CITE-seq data analysis

Raw CITE-seq data from single cells were normalized with the dsb package using the default parameters and empty/background droplets as defined by either clear breaks in the protein library size distribution or droplets defined as negative/background by sample demultiplexing with little impact on normalized values (See Supplementary Note and Supplementary Fig. 9). The denoise.counts argument was set to *TRUE* which carries out the recommended step 2 (denoising cell-cell technical variations by estimating and regressing out the technical component for each cell) and the use.isotype.control argument set to *TRUE* (defining each cell’s technical component by combining isotype control antibody values and the mean of background counts as detailed above). Gaussian Mixture models across cells (as used by the dsb function) or across each protein (as in Supplementary Fig. 1b) were fit using mclust(1). The Uniform Manifold Approximation Projetion(7) (UMAP) Python package was run in R using reticulate with parameters n.neighbors = 35, min.dist = 0.6. Unsupervised protein based clustering was performed using Seurat(8) to implement the SLM(9) algorithm as we previously reported(3) directly on a distance matrix formed on the protein vs cells matrix of CITE-seq proteins (without isotype controls) after normalizing with dsb (in our original report, using pseudocount 1). We retained these cell type annotations used in the original report but renormalized data for all analysis in this paper using dsb with the current package default pseudocount = 10 which resulted in identically distributed relative protein values across cell clusters (Supplementary Fig. 4c see also Fig. 5c in ref 9). In plots of the technical component vs protein library size, nonlinear trends are highlighted by fitting cubic basis splines using the mgcv(10) package implemented with the ggplot function geom_smooth. The full analysis code and data to reproduce all results reported in this manuscript are included in the repository associated with this manuscript (see data availability).

### Assessment of dsb on public external datasets

The raw output from Cell Ranger was downloaded from the 10X Genomics website. Empty droplets and cells were inferred by clear breaks in the distribution of protein library sizes. The same basic steps to normalize protein data that were used for each dataset are illustrated in the dsb package documentation tutorial and vignette https://cran.r-project.org/web/packages/dsb/index.html. The major background/empty droplet population was separated from the cell-containing droplets by an order of magnitude difference in average protein library sizes across droplets. The number of droplets in the upper (cell-containing) peak was consistent with the expected per-lane cell recovery based on the cell loading density of the experiment (Figs. 2g, 2o, Supplementary Figs. 5a, 6a). The droplets in each of these populations were then quality controlled based on mRNA content such that cells must have greater than 200 and less than 3000 unique mRNA and empty droplets must have less than 80 unique mRNA. Cells were then further quality controlled based on the proportion of mitochondrial genes (tuned to each dataset based on density histograms). After these basic quality control steps dsb normalization was carried out using default parameters in the dsb package (denoise.counts = TRUE and use.isotype.control = TRUE). Cells were clustered on a cells by proteins Euclidean distance matrix of dsb normalized values not including isotype control proteins as described above. UMAP was run with n_neighbors parameter = 40 and min_dist parameter = 0.4. Cells are annotated with cluster labels from graph based clustering in Seurat with resolution = 0.8 or 1.2 depending on the dataset and k parameter = 40. The example Mission Bio example data was downloaded from the company’s website. Since this dataset only analyzed ten surface proteins and did not use isotype controls, we performed ambient noise removal, rescaling based on counts in the observed empty droplets only (i.e. performing step 1 only by setting the denoise.counts argument to *FALSE).* UMAP was run with the min_dist parameter set to 0.4 and the n_neighbors argument set to 40 directly on dsb normalized protein values. Clustering was done on a Euclidean distance matrix using Seurat with a resolution parameter set to 0.5 as described above.

### Data/ Code availability

The dsb software is available for download on CRAN: https://cran.r-proiect.org/web/packages/dsb/index.html and is hosted at https://github.com/niaid/dsb. All code needed to reproduce the analysis results reported in this manuscript are available for download at the repository: https://github.com/niaid/dsb_manuscript/. All datasets are all available to download in analysis-ready format for use with the repository above at: https://doi.org/10.35092/yhjc.13370915. The public datasets included in the data repository were downloaded online and are also available from 10X genomics at https://support.10xgenomics.com/single-cell-gene-expression/datasets and from Mission Bio at https://missionbio.com/capabilities/dna-protein/#Data.

## Supplementary Materials

### Supplementary Note

#### Robustness of protein-specific noise estimation assessed by using different definitions of empty/background droplets

The dsb package utilizes the raw (unfiltered) output of UMI count aligners such as Cell Ranger, Kallisto^1^ or as we used here, CITE-seq Count^2^. The unfiltered output (for example, in Cell Ranger, the *raw* output) of droplet barcodes versus UMI counts includes all cell containing and empty (or “background”) droplets, both of which can be inferred using thresholding methods based on the mRNA and protein library sizes-see dsb package documentation tutorial step I for additional details. In all datasets analyzed by us to date, a considerable number (at least 50,000 after QC) of background droplets (i.e., barcodes inferred to not contain at least one cell) can be found using library size based thresholding (see below for robustness assessments). The protein counts derived from these background droplets reflect contributions from ambient antibodies, which as shown in the main text, were highly correlated with the protein counts detected in unstained control cells included in our experiment. Thus, as discussed in the main text, protein counts in empty droplets can serve as an estimate of the expected ambient levels of antibodies. To assess the robustness of estimating protein-specific noise in relation to how background droplets are defined, we compared three approaches to define background droplets. As detailed in our previous report^3^, due to the number of samples included in our experiment, demultiplexing samples required data from both sample barcode (“cell hashing”) antibodies and mRNA (for genetic based demultiplexing, i.e., by cross referencing independently generated patient genotype data using demuxlet, see Methods and main text reference 9). After removing doublets and defining singlets on the basis of data from both the hashing antibodies and genotypes, the remaining (non-doublet, non-singlet) droplets were used to define background droplets in two different ways. First, “Library size background droplets” were defined solely based on library size information where we used clear breaks in the distribution of protein library sizes across the remaining droplets (e.g., as in Figs. 2g and 2o), followed by removal of droplets in the top 10^th^ percentile based on the mRNA library size in order to eliminate droplets containing low quality cells. The library size approach to define background droplets is most compatible with experiments that do not have sample multiplexing, hashing antibody data, such as the external datasets used in this paper (Fig. 2g-v, Supplementary Figs. 5–7).

The second background droplet inference method we tested requires CITE-seq experimental workflows similar to ours, where many samples are multiplexed in the same experiment using sample barcoding antibodies (and/or genetic based demultiplexing). After using Seurat’s K-medoids function to computationally classify cell barcodes as containing singlets, doublets, or negatives based on the hashing antibody counts, we defined “Hashing background droplets” as those classified as “negative” by this demultiplexing software. These droplets had staining below the threshold to be called positive for any one of the hashing antibodies and therefore in principle, their antibody counts should reflect only ambient capture. Such hashing “negatives” were an order of magnitude fewer in number than those determined by library size above, largely due to the threshold used for determining whether a droplet is included in the hash demultiplexing pipeline (the top 35,000 barcodes from each lane). Hashing background droplets were further filtered to: 1) include only droplets classified as “ambiguous” by SNP demultiplexing (via demuxlet), i.e., these cannot be attributed to a single or multiple distinct donors based on cross-referencing mRNA reads in the droplet with independently generated genotype data, and 2) exclude any droplet with >80 unique mRNAs to remove cell-containing droplets with low-quality mRNA capture. Using this alternative method to define background droplets, we similarly observed that the relative amount of antibody was highly correlated between unstained cells and these background droplets (along the unity line in Supplementary Fig. 1b).

Interestingly, while the correlation was similarly high, antibody levels in unstained cells or in hashing background droplets were greater than those in library size background droplets (Supplementary Fig. 1b top vs bottom). The greater magnitude of antibody counts by a multiplicative factor in log-count space (slope in bottom panel of Supplementary Fig. 1b is 1.24 with near zero intercept) suggests that unstained cells and demultiplexing background droplets capture additional antibodies. Unstained cells may serve as an additional antibody capturing “reservoir”, e.g., due to non-specific (or specific) binding. However, this would not explain their concordance with demultiplexing background, which, as supported by both genetic (via demuxlet classification) and barcoding antibody (via Seurat k-medoids classification) data, should have a low chance of containing fully intact cells. It is still possible, despite filtering out droplets with low mRNA counts, that demultiplexing background droplets contained some very low-quality cells or cell membrane debris that together could capture additional antibodies from the environment via specific/non-specific binding. Demultiplexing background droplets could also have more ambient mRNA (as described above in order to be included in the hashing antibody demultiplexing step) than droplets defined using the protein library size distribution alone, and thus they (as also in the unstained control cell droplets) could conceivably serve as an additional set of free antibody-capturing molecules. Importantly, however, we emphasize the difference between empty/background droplets defined using protein library size distribution versus hashing antibody demultiplexing had negligible effect in the resulting dsb normalized values (see below).

We further investigated a third approach to estimate protein background noise–the mean of each protein across the subset of *stained cells* that were inferred to belong to the *“negative”* population for each protein. Without dsb rescaling, we fit a two component Gaussian mixture model to the log + 1 transformed count of each protein *i* across single cells, resulting in 2 populations of cells: those positive or negative for the protein. Each protein’s background mean, *“A”* (see Supplementary Fig 1a), reflects the average log transformed count of the non-staining cell population for that protein, i.e., cells that do not express that protein. The protein level in unstained controls and empty drops were both highly correlated with *A* (Supplementary Fig. 1c). Thus, antibody levels in unstained droplets on average are similar to those in droplets with stained cells not expressing the target protein.

We thus have tested three different ways to estimate the average background protein noise correlated across droplets. We further found that the noise signal captured in library size background droplets appears to be universally found in data from all of the droplet-based oligo barcoded antibody experiments we examined and is thus a generalizable method of estimating noise (e.g., see Fig. 2g, 2o and Supplementary Figs. 5a, 6a, 7a). A recent study evaluating experimental strategies to improve signal to noise in CITE-seq experiments used the dsb package to normalize CITE-seq data from different titration experiments^4^. Consistent with our finding that ambient antibodies are major contributors to the correlated noise component across single cells, this report found that background noise was correlated with staining concentration, i.e., the highest background noise was found for antibodies with higher staining concentration (10μg) and low in the estimated cognate antigen density on cells available to specifically bind the antibody^4^.

#### Importance of using isotype control antibodies for estimating cell-intrinsic normalization factors

Despite μ1 alone providing information about the cell-intrinsic technical component, we recommend the inclusion of multiple isotype controls in CITE-seq experiments to serve as anchors for revealing technical normalization factors because μ1 alone may carry signals beyond those from technical factors (e.g., low-level antigen specific binding). In our data for example, μ1 exhibited greater correlation with μ2 than did the mean of the isotype controls (Supplementary Fig 3b,c), even when sub-sampling random draws of four proteins from those used to compute μ1 within each cell to assess whether signal from four background proteins is equivalent to that of four isotype controls (Supplementary Fig. 3d). Furthermore, even with isotype controls as anchors, the estimated cell-intrinsic background may encompass signals from non-specific binding to surface Fc receptors. Cell types such as monocytes with higher relative Fc receptor expression may thus receive more correction than other cell types. However, empirically we have not found this to have adverse effects on normalized values in populations such as monocytes, cell type identification, or downstream analysis (Supplementary Figs. 4b, c). Careful blocking of Fc receptors before antibody staining, which is standard practice and was performed in our experiments, likely contributed to mitigating this effect.

#### Robustness of dsb normalized values to background droplet definition and multi-vs single-batch processing

Given the strong correlation observed between average protein levels in unstained control cells and droplets with stained cells expressing background level of the protein (Supplementary Fig. 1c), ambient antibodies appear to capture the major noise component that contributes to each protein’s specific noise floor. In external datasets, we defined background droplets by using thresholds determined based on the protein library size for each dataset and excluded droplets with more than 80 unique mRNA. This procedure revealed a clear population of more than 50,000 background droplets in each dataset. In some external datasets, there were two distinct background populations based on protein library size, which had library sizes an order of magnitude lower than the positive, cell-containing droplets (Supplementary Fig. 8a). dsb normalized values were robust to using different background subpopulations (Supplementary Fig. 8b). When only the lower background peak was used to simulate an experiment with extremely low background, dsb counts still separated canonical cell populations but were less 0-centered due to the low estimated background for some proteins (third row, Supplementary Fig. 8b). However, this is only a theoretical limitation as we have not encountered such a dataset to date.

The experimental design of the main dataset used here to develop our approach include n=20 unique donors distributed over two experimental batches; this presented multiple options for dsb normalization. Background/empty drops could be defined with either of the two methods described above (demultiplexing or library size), and cells could then be normalized by combining all cells / background into a single matrix and normalizing both batches together, or each batch of cells could be normalized separately, using only the empty droplets within each batch. To test how robust the resulting dsb normalized values were to single vs multi-batch normalization, as well as to further validate the findings described above on the robustness of dsb normalized values to different definitions of background, we tested the 4 possible normalization schemes with background droplets classified by either protein library size distribution or demultiplexing then normalized with dsb by either merging cells and background from both batches together, or normalizing each batch separately. The resulting dsb normalized values were consistently similar across all four of these normalization schemes (Supplementary Fig. 9). Since we expect ambient antibody to be a major contributor of correlated noise across cells, experimental standardization of staining time and the number of washing steps prior to droplet generation could be important contributing factors in mitigating batch to batch variations. In our dataset, both batches were run by the same experimenters on subsequent days with the same pool of antibodies using the exact same protocol, which may have contributed to the lack of major batch to batch variability.

---

**Supplementary Figure 1.**
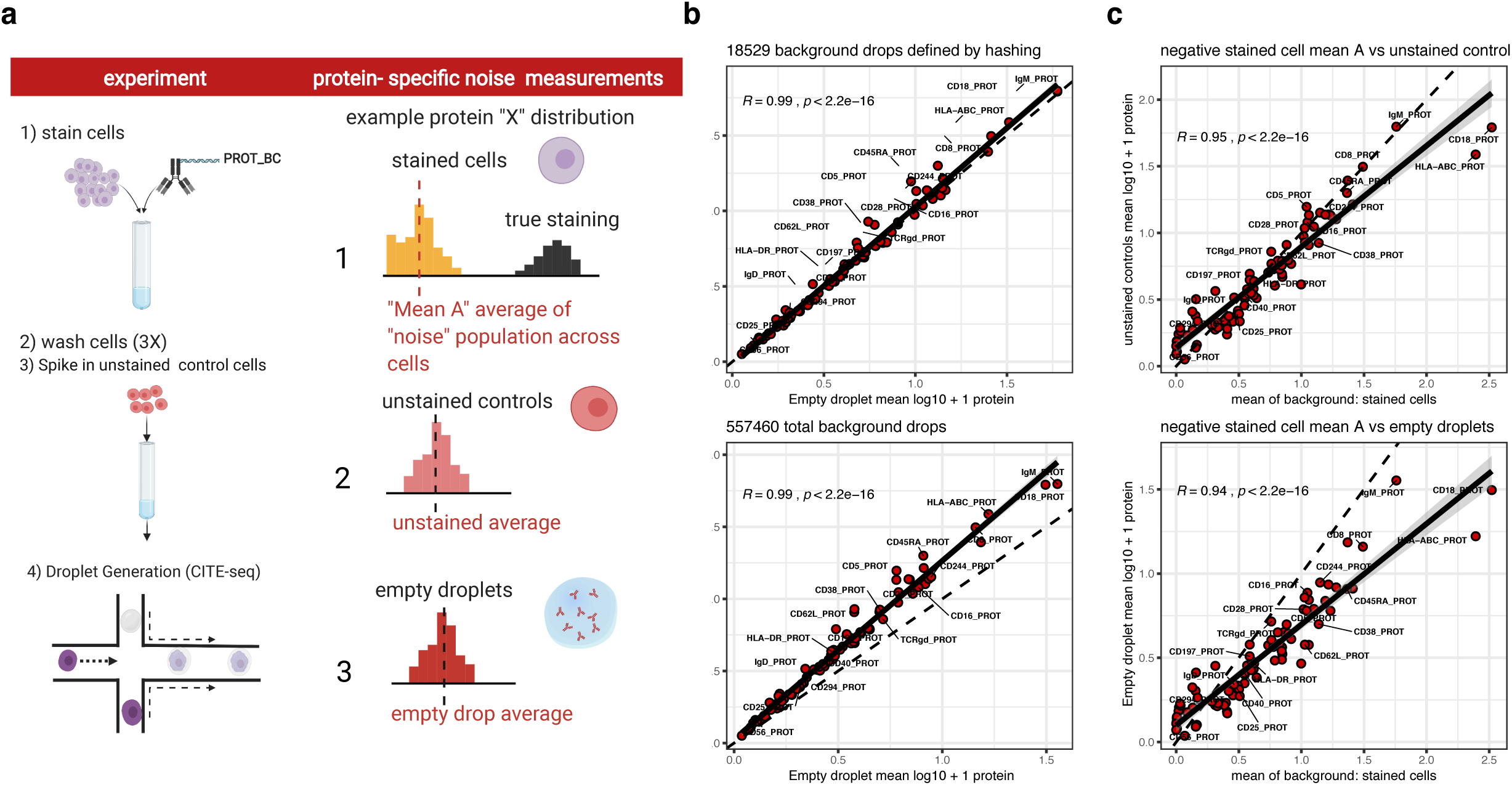
**a.** Expanded from Fig. 1a: to assess the relative contribution of the ambient antibody component of noise correlated across droplets, three different measurements of protein specific background noise were defined for each protein: 1) (top row, right column) for each protein, the average log transformed value of the subset of stained cells that were not part of the proteins “positive” population and comprised the “non-staining” population of cells (the negative cell population for each protein was inferred through a Gaussian mixture model fit separately to each protein, see Methods) 2) (middle row, right column): unstained control cells spiked into the cell mixture prior to droplet generation as shown in the experiment diagram (left column), 3) (bottom row, right column): empty droplets as defined by either the protein library size distribution or inferred by sample barcode antibody demultiplexing (see Methods). **b.** Pearson correlation between unstained control cells (y-axis) and empty droplets (x-axis) with empty droplets defined by either demultiplexing (top “hashing background droplets”) or library size distribution (bottom, “library size background droplets”, see supplemental note) **c.** Pearson correlation between y-axis: unstained controls (top panel) or library size background droplets (bottom panel) versus x-axis: the mean of the protein in stained cells that were negative for the protein (“mean A” as shown in top panel of a). In all plots the dashed line at unity (y = x) is shown for reference and the solid line is the fitted regression line with the shaded region representing the 95% confidence interval of the linear model fit.

**Supplementary Figure 2.**
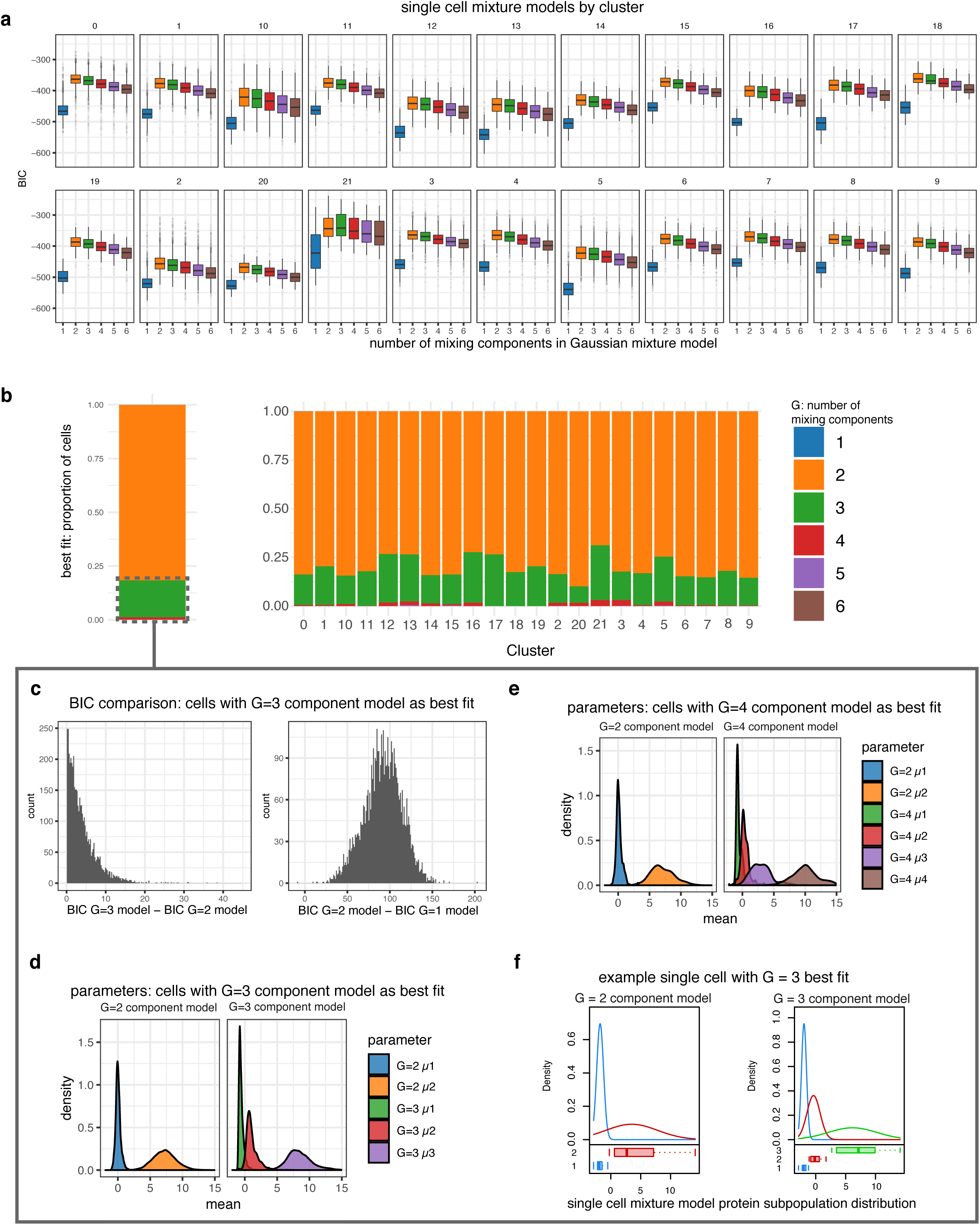
Assessment of the modeling assumptions for defining each cell’s background protein population mean *μ1* with a k=2 component mixture model for use in the per-cell technical component regressed out of dsb normalized counts in step II–see related figures on external validation datasets Fig. 2 k, s, Supplementary Figs. 5,6h. **a.** Gaussian mixture model fits (from (a) and Fig. 1d) partitioned by each protein-based cell cluster (clusters are the same as defined after dsb normalization in Kotliarov *et. al.* 2020). **b.** In more than 80% of cells (shown in orange, left bar plot panel) the Gaussian mixture model with two subpopulations maximizes the BIC. In the 17% of cells where G = 2 was not the best fit, the G = 3 model had the best fit (shown in green), these cells with G = 3 as the best fit were not biased to a specific protein-based cluster (bar plot for each cluster on right). **c.** For the cells with G = 3 models having the best fit, the distribution of the difference in BIC between G = 3 vs. G = 2 (left) and G = 2 vs. G = 1 (right) models shows the fits for G=2 vs 3 are similar, whereas the single mean model had a comparably much lower BIC (lower BIC corresponding to models with a worse fit using conventions from the mclust package-see Methods) than the two-component model. **d.** The distribution of Gaussian mixture model subpopulation means for G = 2 and G = 3 models for the subset of cells with G = 3 as the optimal fit shows G = 3 and G = 2 models fit similar values for μ1 in these cells. **e.** As in (d); the small minority of cells (shown in red in (c)) with G = 4 as the best fit. **f.** A single arbitrary example cell that had an optimal BIC with the G = 3 model; the distribution of inferred mixture model means is shown for the 2-subpopulation (left) and 3-subpopulation (right) model fits showing overlapping value for *μ1*.

**Supplementary Figure 3.**
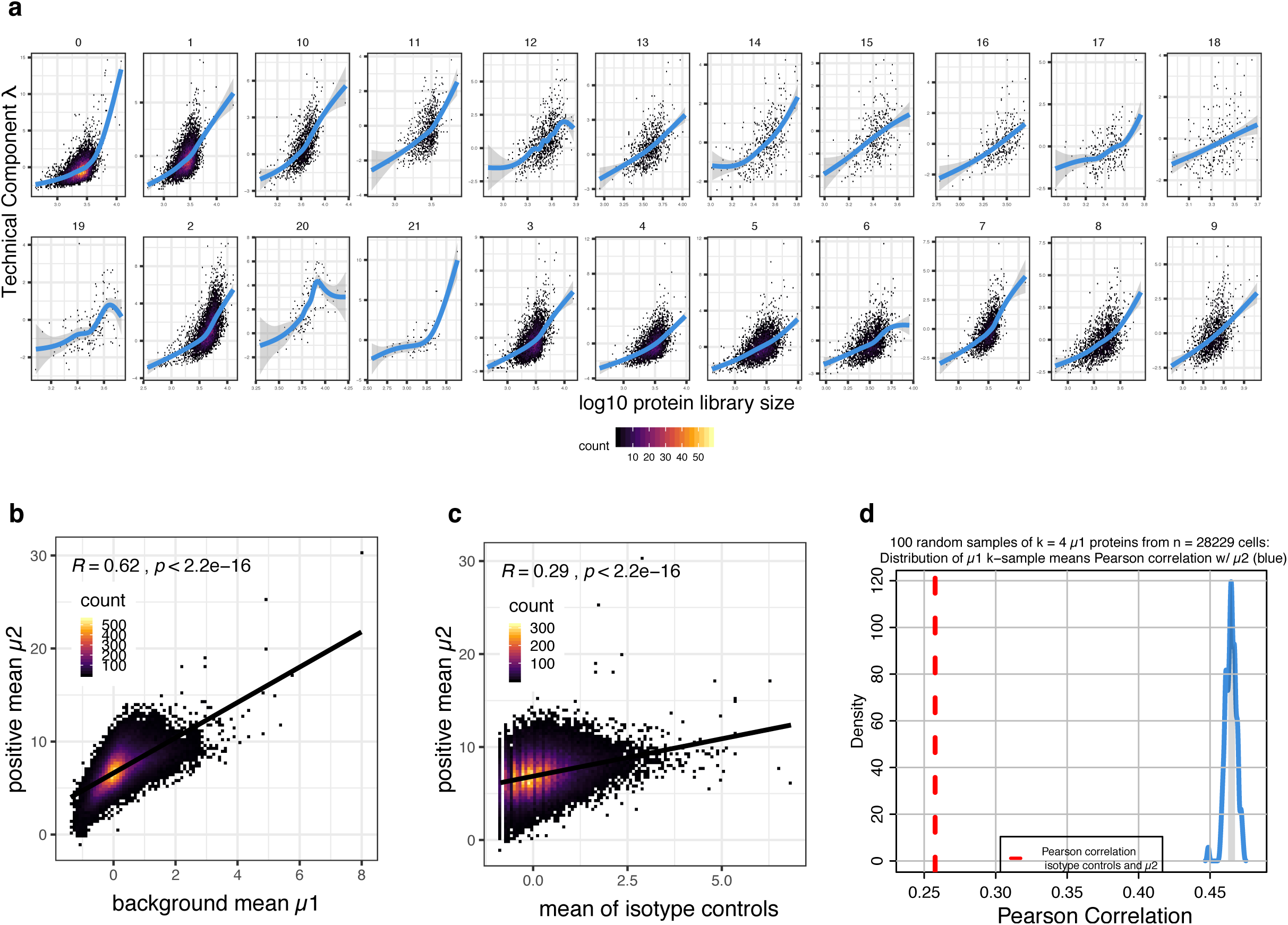
**a.** As in Fig. 1g, each cell’s inferred technical component λ (y-axis) defined as the first principal component through the isotype controls and the background mean (blue distribution, Fig. 1c, see Methods) vs the cell’s protein library size (defined as the sum of the protein UMI counts in each cell). Each number indicates protein-based clusters (see Methods) as shown in Supplementary Fig. 4. The nonlinear trend is highlighted with a cubic basis spline. **b.** The Pearson correlation between μ1 and μ2 from single cell k = 2 component mixture models fit across all proteins in each cell. **c.** The average of isotype controls after dsb normalization step I (ambient correction) vs μ2 as in (b). **d**. The distribution of n=100 Pearson correlation coefficients between each cell’s μ2 and 100 random samples of k=4 μ1 proteins from each single cell (blue) shaded region is the 50% highest density interval, red line is the Pearson correlation coefficient of μ2 and the mean of isotype controls in each cell.

**Supplementary Figure 4.**
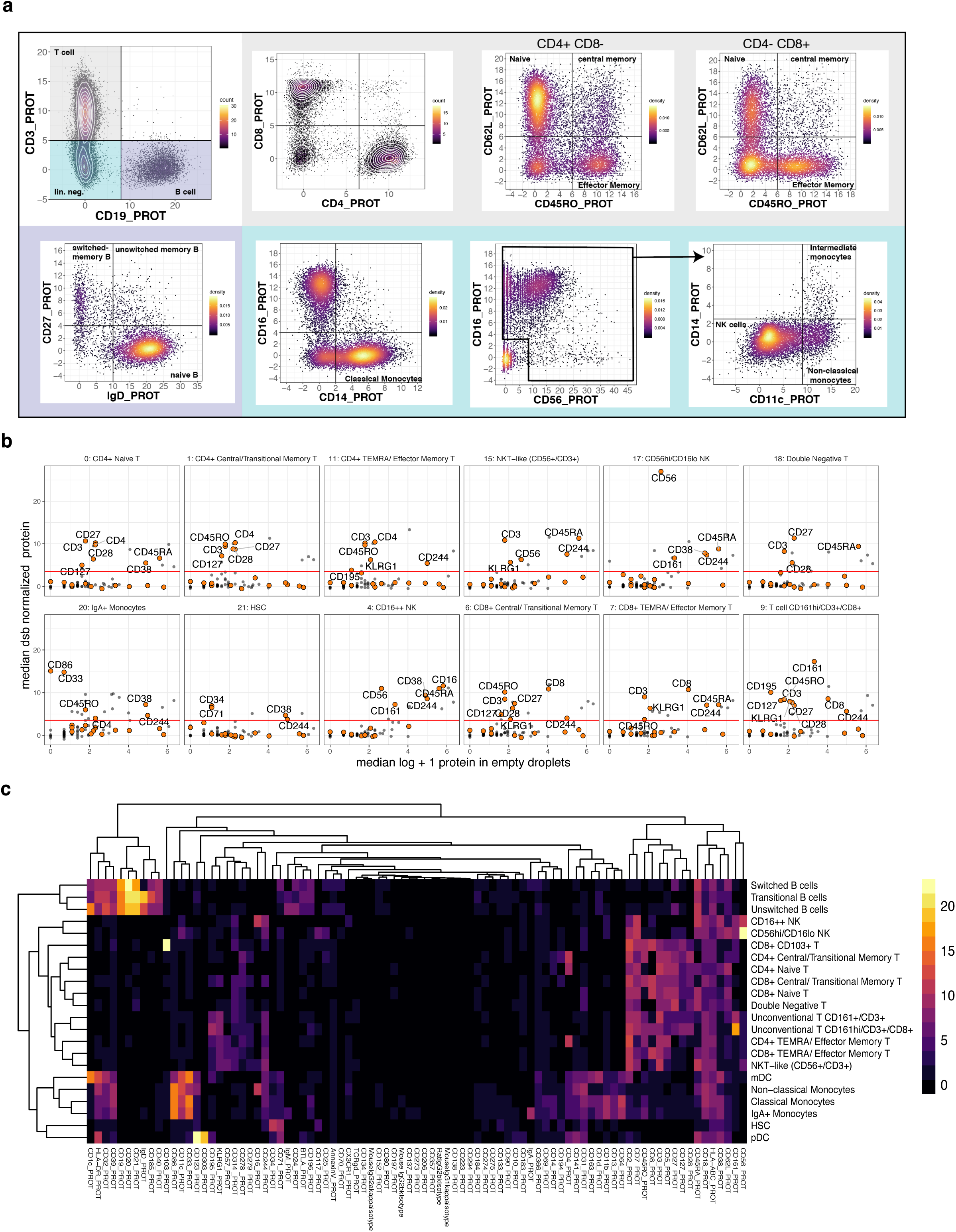
**a.** Biaxial gating strategy for identifying major immune cell subsets with dsb normalized values. Grey = T cells, Blue = Monocytes, Purple = B cells. **b.** As in Fig. 2f, the average log transformed protein count in empty droplets (x-axis) vs the average dsb normalized values (y-axis) for each protein-based cell cluster–the threshold above which proteins are annotated in the plot is 3.5 corresponding to 3.5 standard deviations above expected noise +/- the technical component correction applied in step II (see methods). The subset of proteins highlighted in orange in these panels are CD56, CD71, CD27, CD244, KLRG1, CD195, CD38, CD127, CD16, CD3, CD4, CD8, CD28, CD161, CD45RO, CD45RA, CD34, CD33, and CD86. **c.** Heatmap of average dsb protein normalized expression in each cluster.

**Supplementary Figure 5.**
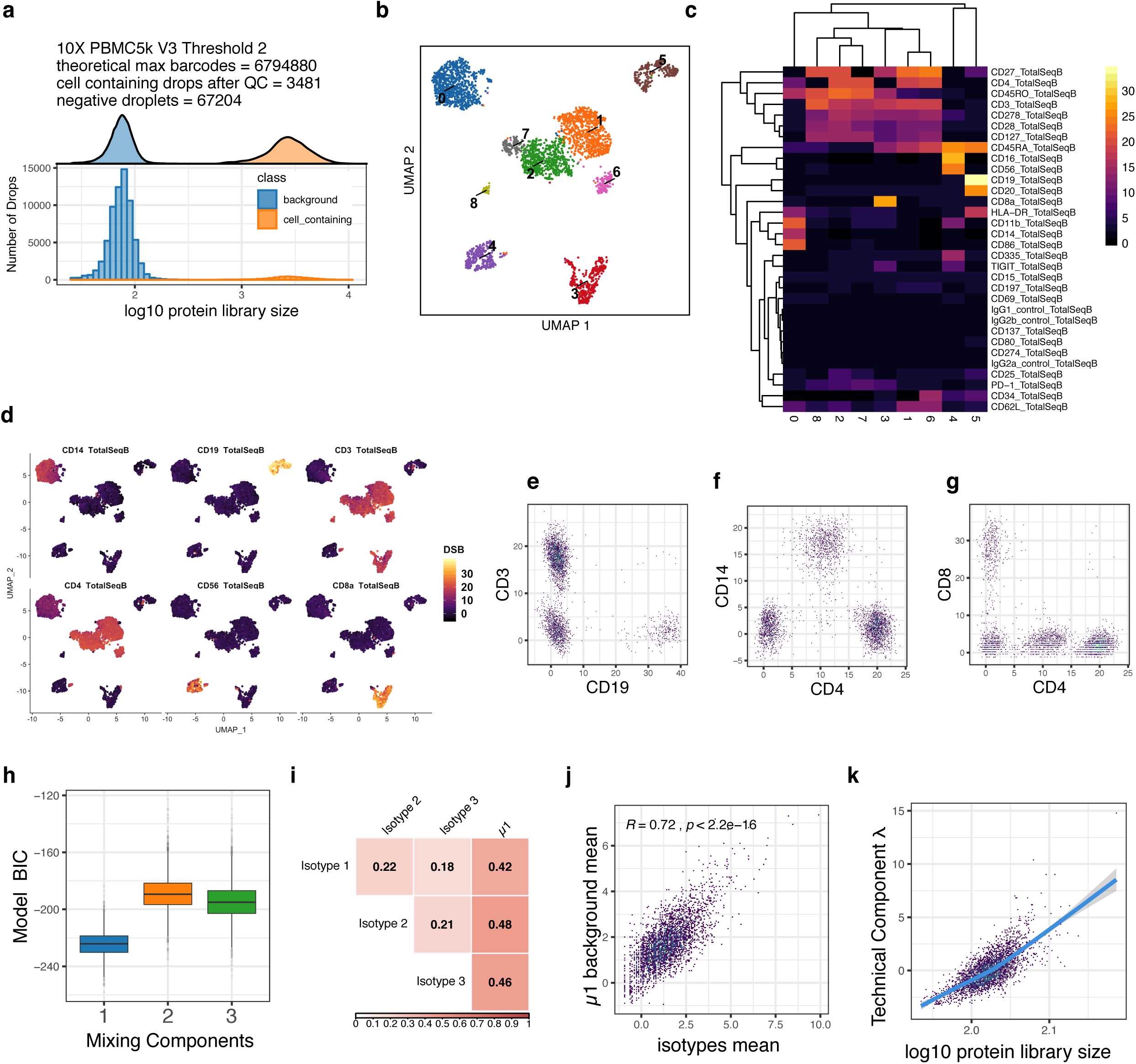
Data from 10X Genomics Chromium “5 Prime” assay from the “5’ 5K PBMC dataset”. **a.** As in Fig. 2g, o: the protein library size distribution of empty drops and cell containing drops used for dsb normalization. **b.** As in Fig. 2h,p UMAP of cells clustered by dsb normalized surface protein. **c**. as in Fig. 2i,q, heatmap of the average of dsb normalized values per protein-based cluster shown in (b). **d.** As in Fig. 2e, a UMAP plot with cells labeled by dsb normalized protein expression for canonical immune cell markers. **e-g.** As in Fig. 2j,r biaxial plots show canonical immune cell populations after dsb normalization. **h.** As in Fig. 2k,s the Gaussian mixture model fit to the dsb normalized values of each single cell from step I (ambient correction). The model Bayesian Information Criterion (BIC) vs mixing components in the model is shown with G= 2 models globally having the best fit. **i.** As in Fig. 2l,t, correlation matrix of variables used to define each cell’s technical component; each isotype control and μ1, the Gaussian mixture model background mean across proteins for each cell. **j.** As in Fig. 2m,u, Pearson correlation coefficient between the inferred cell-specific background mean μ1 from the Gaussian mixture model vs the mean of isotype controls in each cell. **k.** As in Fig. 2n,v, the relationship between each cell’s technical component and the cell’s protein library size.

**Supplementary Figure 6.**
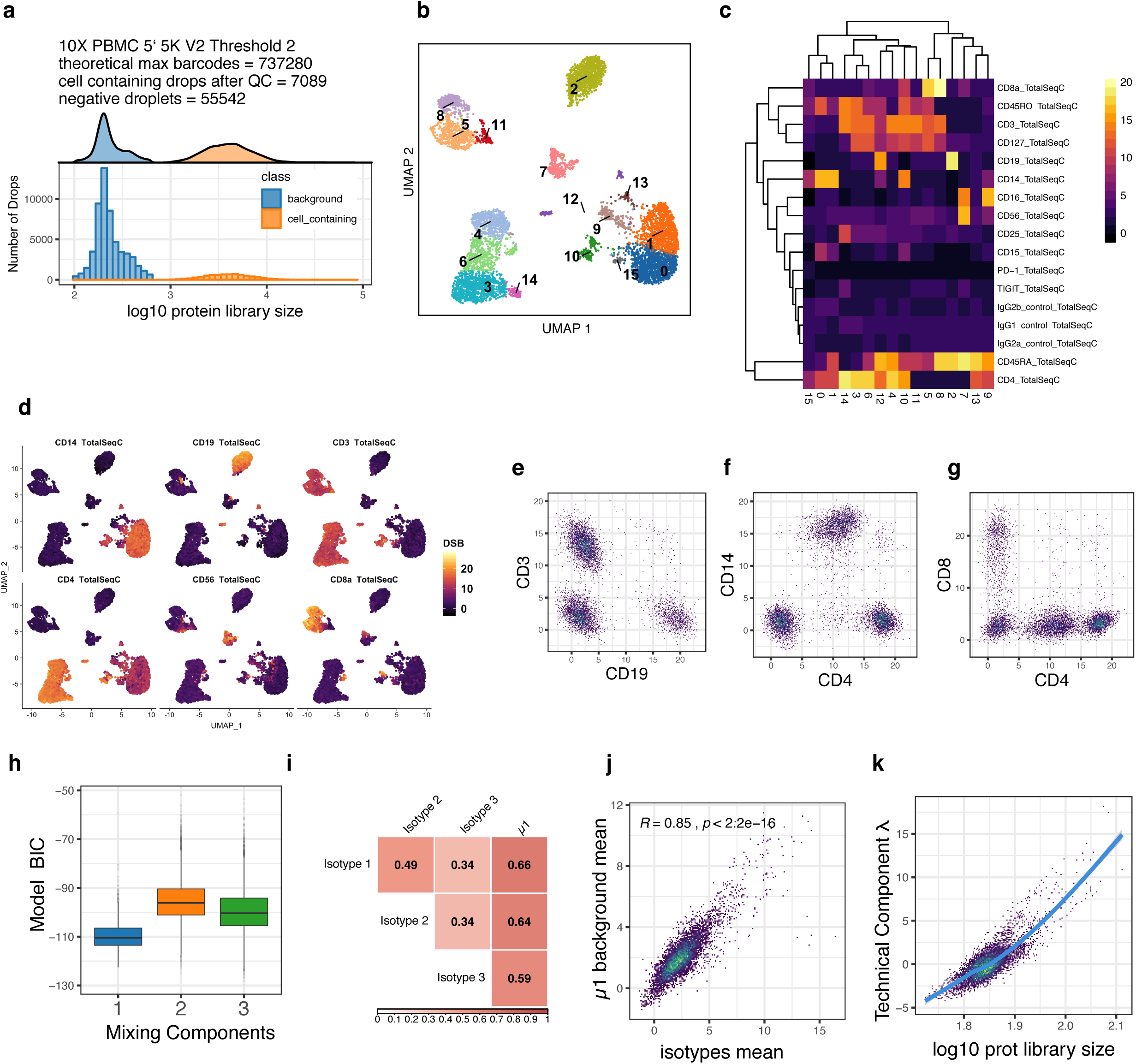
Similar to Supplementary Figure 5, showing analysis of data from 10X Genomics Chromium Version 3 “PBMC 10K data”.

**Supplementary Figure 7.**
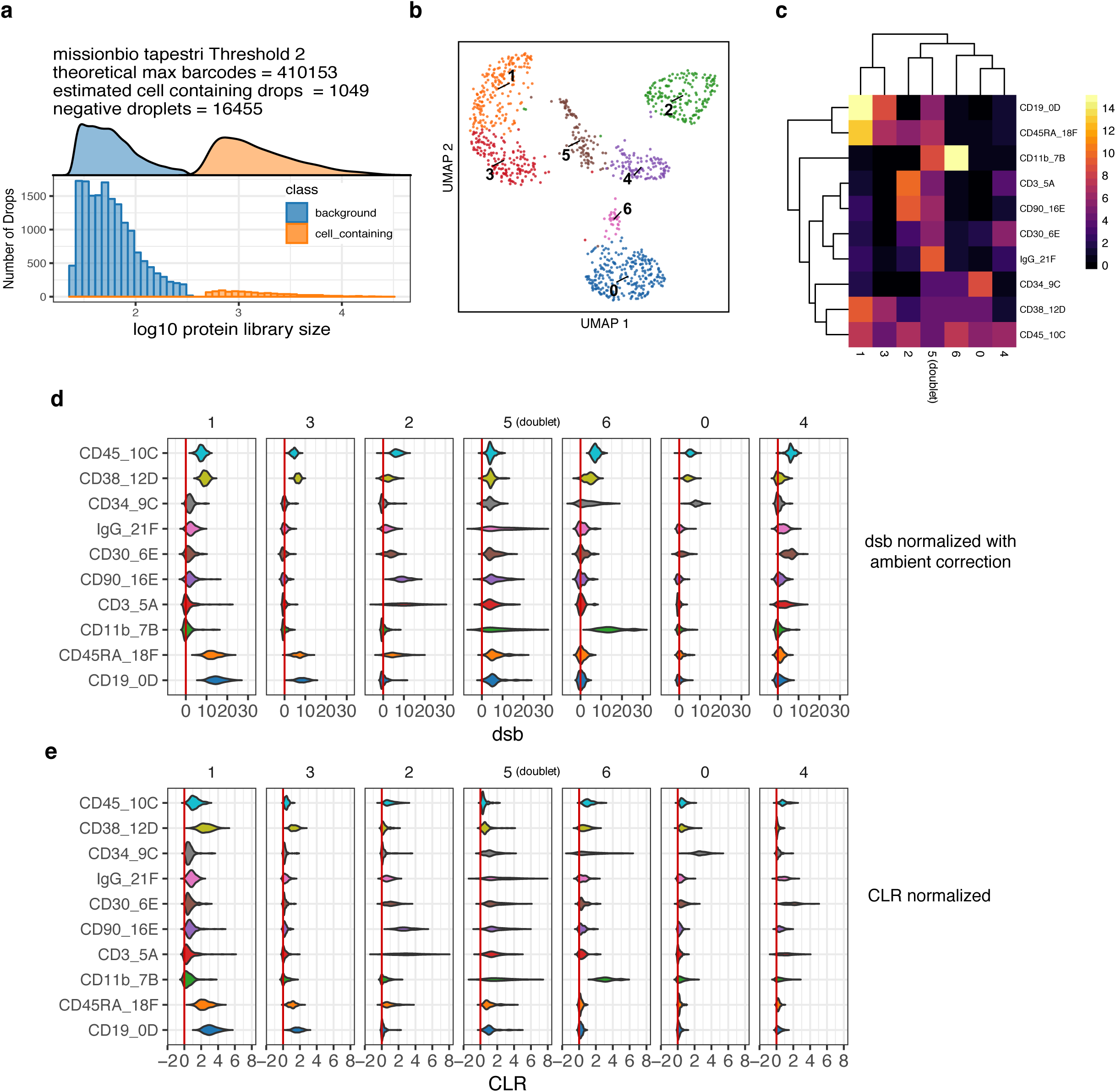
A mixture of n=4 leukemia cell lines from example data for the Mission Bio ‘Tapestri” platform for simultaneous “proteogenomic” assessment of surface proteins and DNA **a.** The protein library size (total UMI) distribution was used to estimate cell-containing droplets and empty droplets without a cell, as in the analysis of 10X Genomics data using the same procedure shown in Fig. 2g,o and Supplementary Fig. 5a and 6a. **b.** UMAP analysis based on dsb normalized values, cells are labeled by graph-based cluster identity. **c.** heatmap of the average expression of each dsb-normalized protein in each cluster. The range of values is on the same scale for all proteins ranging from less than 0 to 14, corresponding to 14 standard deviations from the background observed in empty droplets–cell to cell technical variations were not inferred through calculating the technical component for each cell in this dataset due to the number of proteins profiled (n=10) and the lack of isotype controls from which to define each cell’s technical component (see Supplementary Note and Methods). **d.** Violin plots for protein expression within each cluster as normalized with dsb or with **e.** CLR transformation.

**Supplementary Figure 8.**
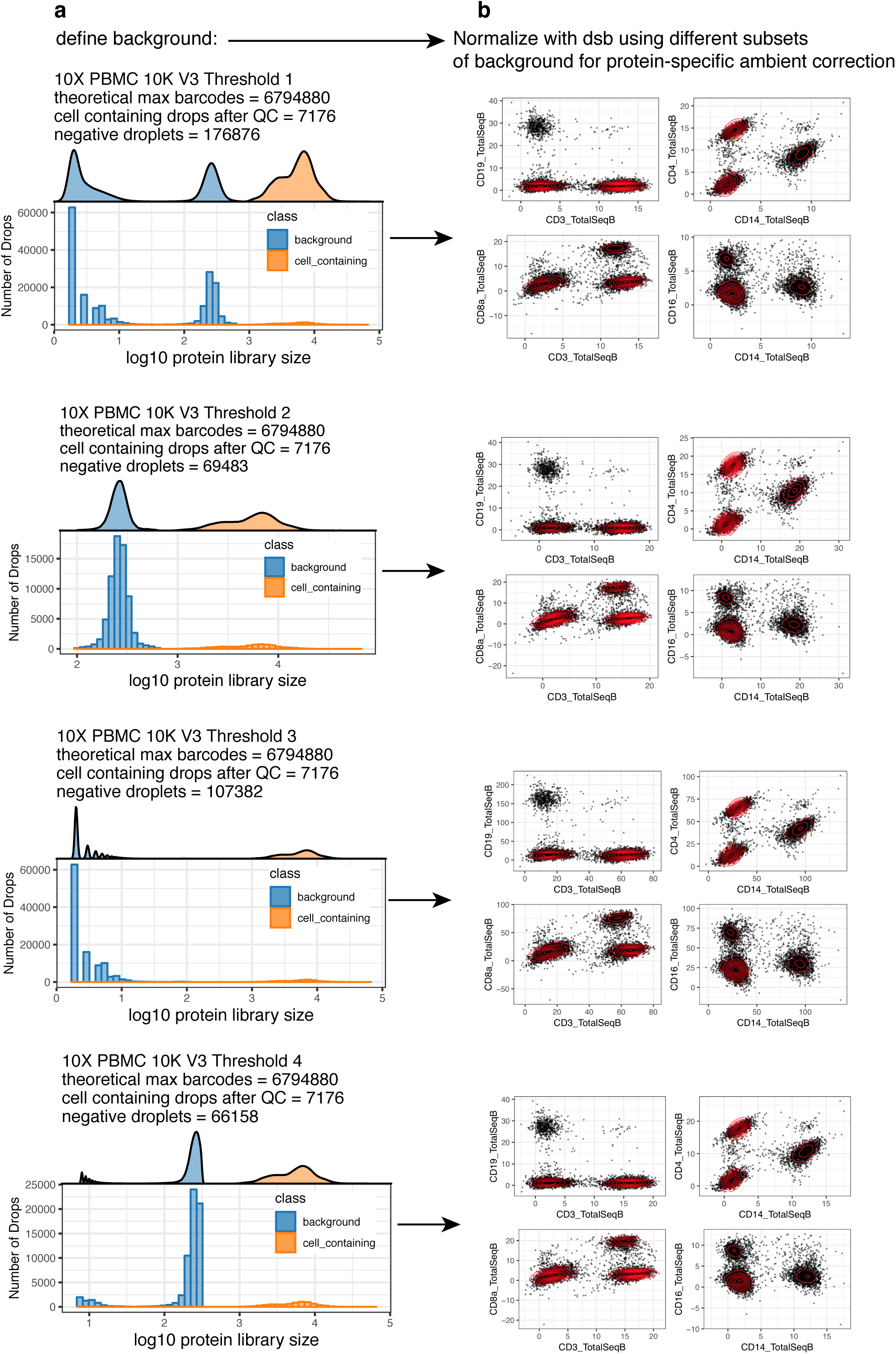
Robustness of dsb normalized values to different definitions of background droplets. **a.** Distribution of protein library size for the 10X genomics Chromium Version 3 “PBMC 10K” dataset which had a bimodal distribution for the non-cell-containing droplets shown in blue. In each row, a different threshold based on the protein library size was used to define background droplets, which were then used to normalize the same population of cell-containing droplets (shown as the orange distribution) with the dsb package. **b.** The dsb normalized values are shown for canonical protein-based phenotypes with biaxial scatterplots. The scale of the 3^rd^ row is negatively impacted by eliminating the major empty droplet background peak with greater mean value and only using the empty droplet background peak with very low mean library size.

**Supplementary Figure 9.**
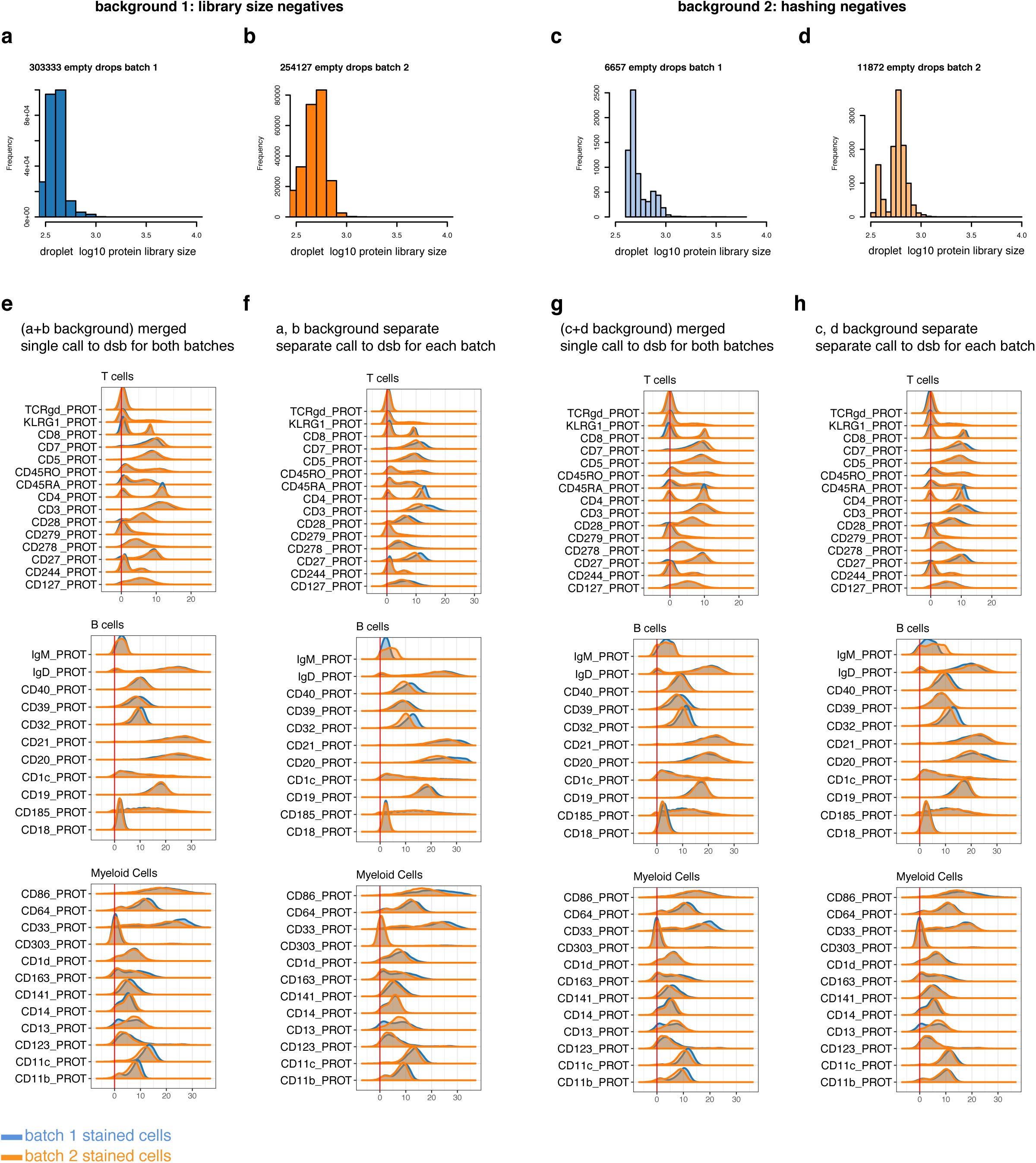
Stability of dsb normalized values when processing multiple batches in a single normalization vs normalizing each batch separately, both using two definitions of background droplets with the dsb package. **a-d** show protein library size distributions of background droplets defined using either the protein library size distribution alone or droplets defined as negative during demultiplexing (see Supplemental note) across n = 2 batches. The raw Cell Ranger outputs from each staining batch of cells were split across n=6 lanes of the 10X Chromium instrument and for each definition of background, the dsb results for merged vs split batch normalization are shown. **e.** Merged normalization: empty drops from both batches and the cell containing droplets for both batches are each combined into a single matrix of cell-containing and empty drops and normalized with dsb. **f**. Split batch: empty drops and cell-containing drops from each batch are used in two separate normalizations with the dsb package with normalized values combined into a single matrix post normalization. The distributions shown in **g** and **h** are computed identically to **e** and **f** (**g** = merged batch, **h** = split batch) and the only difference is the empty droplets used to define the background (distributions shown in **c** and **d** – the “hashing background droplets” see Supplementary Note).

## Supplementary software documentation

This is a snapshot of select topics covered i n the dsb vignette hosetd on CRAN. For the current version of the ful l software vignette please see: https://cran.r-project.org/web/packages/dsb/index.html For comments please see https://github.com/niaid/dsb

### Using dsb to normalize single cell protein data: analysis workflow and integration with Seurat, Bioconductor and Scanpy

dsb (**d**enoised and **s**caled by **b**ackground) is a lightweight R package developed in **[John Tsang’s](https://www.niaid.nih.gov/research/john-tsang-phd)** Lab (NIH-NIAID) for removing noise and normalizing protein data from single cell methods such as CITE-seq, REAP-seq, and Mission Bio Tapestri. **[See the dsb Biorxiv preprint](https://www.biorxiv.org/content/10.1101/2020.02.24.963603v1)** for details on the method and please consider citing the paper if you use this package or find the protein noise modeling results useful.

This vignette outlines how to use dsb to normalize droplet-based single cell protein data from antibody counts and how to integrate the normalized values in an analysis workflow using Seurat, as well as integration with Bioconductor and Scanpy. If you have a question please first see the FAQ section then open an issue on the github site https://github.com/niaid/dsb/ so the community can benefit.

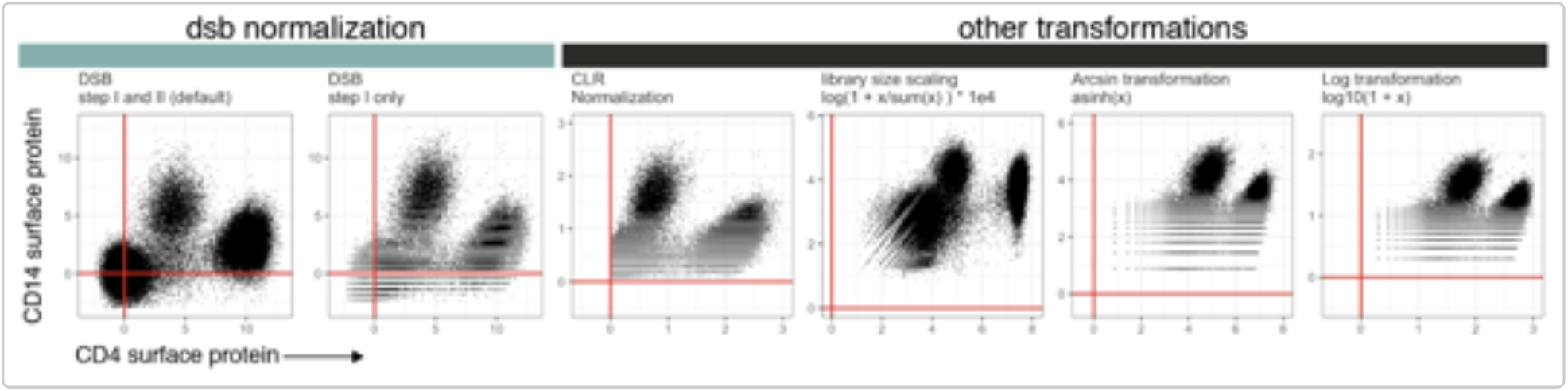

#### Background and motivation for the dsb method

CITE-seq protein data suffers from substantial background noise (for example, see supplementary fig 5a in Stoeckius et. al. 2017 Nat. Methods). We performed experiments and analysis to dissect this noise and the dsb method is based on 3 key findings outlined in our paper.

1. Based on unstained control experiments and modeling we found a major source of protein background noise comes from ambient, unbound antibody encapsulated in droplets.
2. “Empty” droplets (containing ambient mRNA and antibody but no cell), outnumber cell-containing droplets by 10-100 fold and capture the *ambient component* of protein background noise.
3. Cell-to-cell technical variations such as stochastic differences in cell lysis/capture, RT efficiency, sequencing depth, and non-specific antibody binding can be estimated and removed by defining the “technical component” of each cell’s protein library.

#### Installation and quick overview with pre-loaded package data

Use the `DSBNormalizeProtein()` function to

1) normalize raw protein counts of a cells (columns) by proteins (rows) matrix `cells_citeseq_mtx` estimating the ambient component of noise with `empty_drop_citeseq_mtx`, a matrix of background/empty droplets. 2) model and remove the ‘technical component’ of each cell’s protein library by setting `denoise.counts` = TRUE and include isotype controls in the calculation of the technical component with `use.isotype.control` = TRUE.

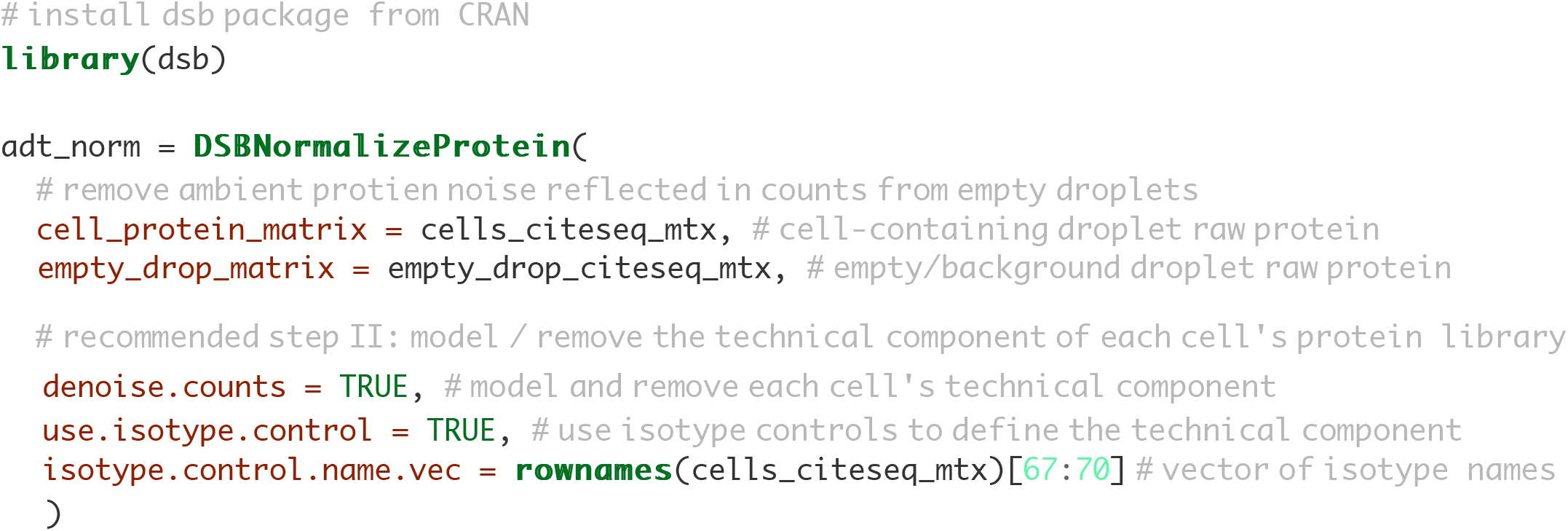

#### Tutorial with public 10X genomics data

Download **RAW (*not* filtered!)** *feature / cellmatrix raw* from public 10X Genomics CITE-seq data here. The tutorial below uses use R 3.6. We emphasize normalized protein data and raw RNA data from this workflow at step III can be used with Seurat, Bioconductor or Python’s AnnData class in Scanpy. We use a convenience function from S eurat Reαdl0X to load the raw data. We then provide a suggested workflow after normalization with dsb based on the CITE-seq analysis used in our paper on baseline immune system states: Kotliarov et. al. 2020 Nat. Medicine.

Please see **multiplexing experiments section** if your experiment used superloading/demultiplexing and **FAQ section** for guidance if you have multiple 10X lanes and or batches.

#### Step Ia: load raw count alignment (e.g. Cell Ranger) output and define cell metadata variables

Below we load the **raw** output from the Cell Ranger count alignment. The raw output is a sparse matrix of **possible cell barcodes** vs proteins / mRNA. The number of cell barcodes ranges 500k-6M depending on the kit/chemistry version.

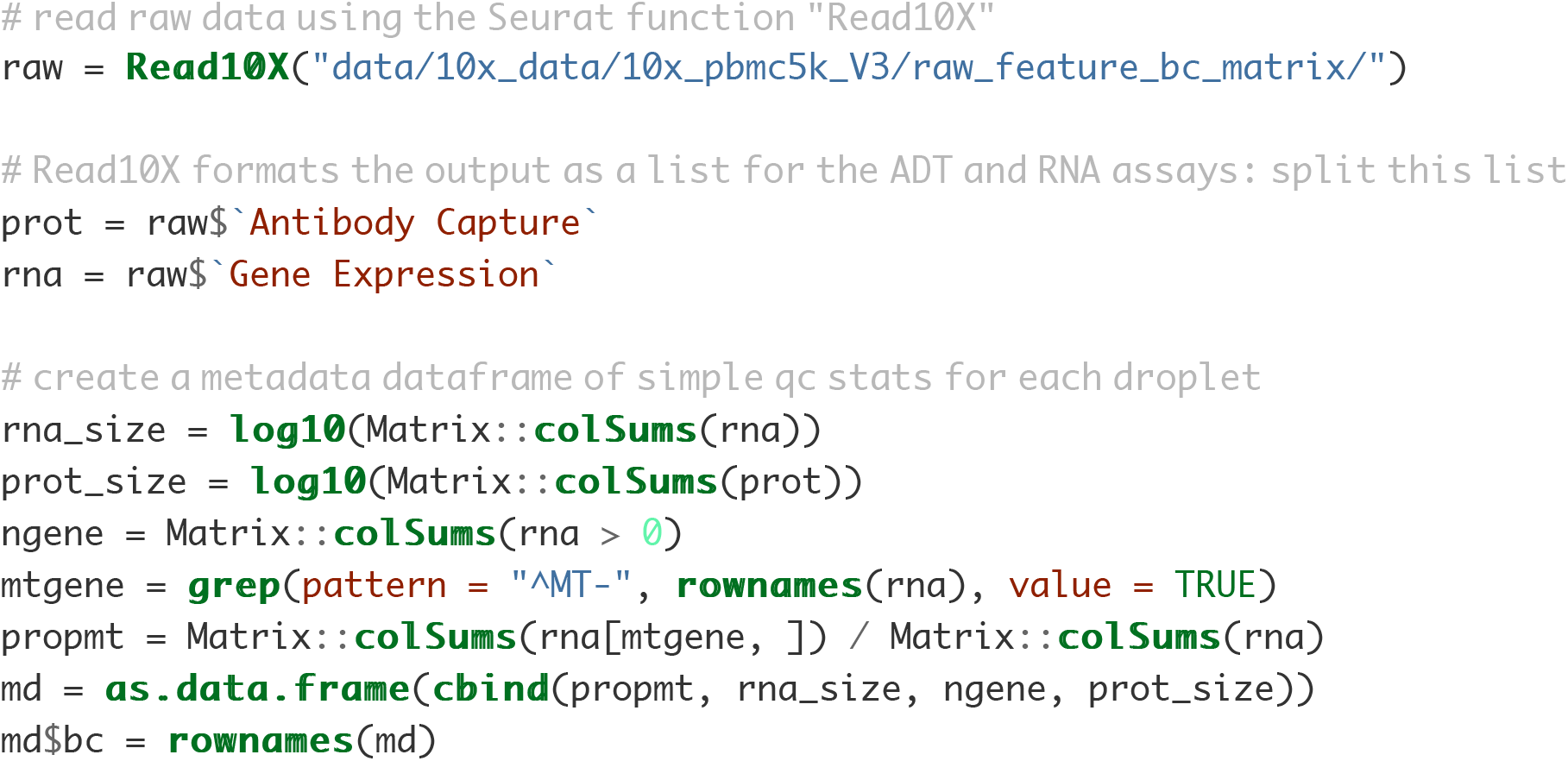

#### Step Ib: Quality control and defining cell-containing and background droplets

Now that we loaded the raw output, we define cells and background droplets. Methods for defining droplets depend on the experiment design. The number of droplets you define as cell-containing should be in line with the number of cells loaded and the expected cell recovery. In this experiment, 5000 cells were loaded in a single 10X lane. In the protein library size histogram below, large peaks < 1 and ~2 contain > 30,000 and 60,000 drops respectively. The smaller peaks > 3 (for both mRNA and protein) contain ~5000 drops and therefore **the rightmost peaks > 3 are the cell-containing droplets** we next perform quality control as in any single cell preprocessing workflow. Again, see multiplexing experiments section and FAQ if you have many lanes / batches and or superloading.

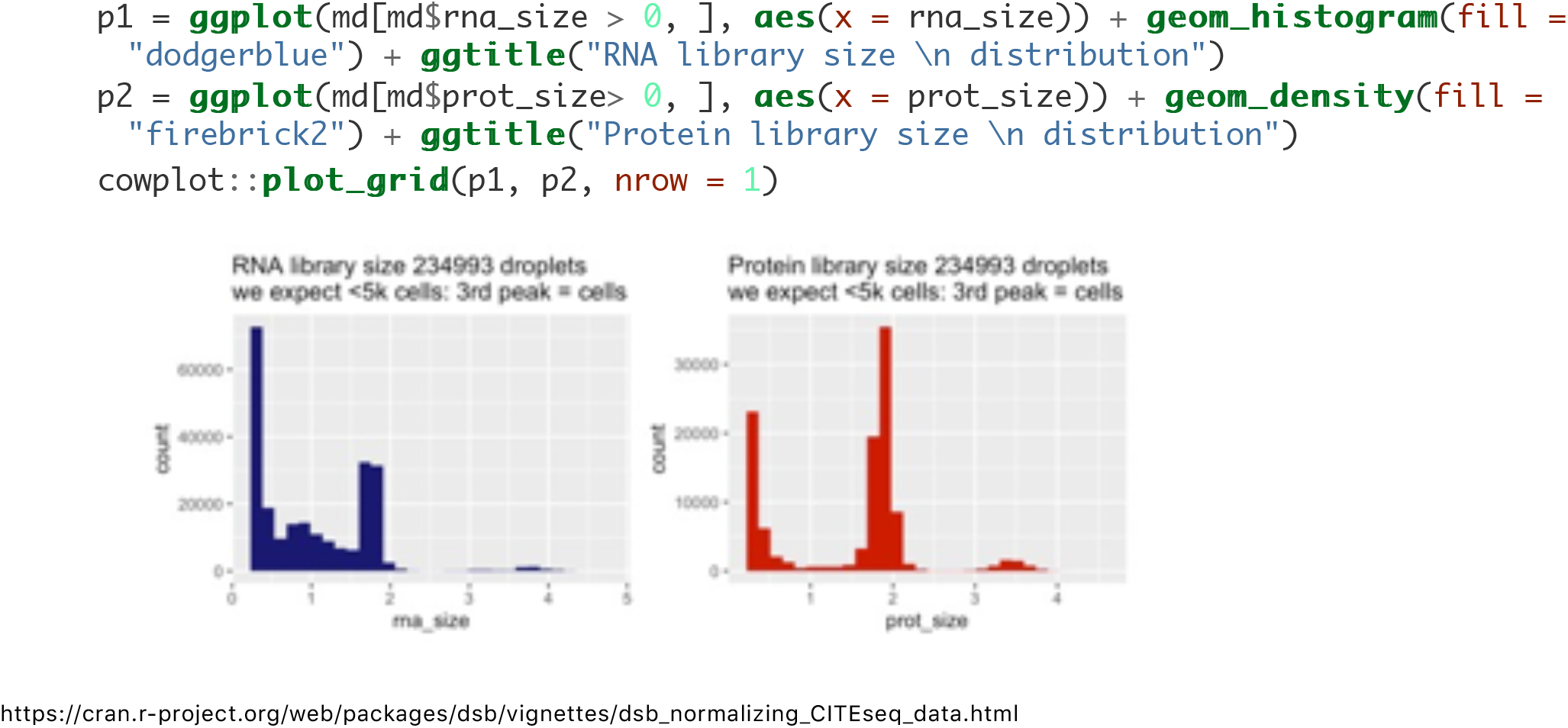

Below we define background droplets as the major peak in the background distribution between 1.4 and 2.5 log total protein counts-note one could also define the empty/ background droplets as the entire population from 0 to 2.5 with little impact on normalized values (see the dsb paper section on sensitivity analysis to different definitions of background for details). In addition, we add mRNA quality control-based filters to remove potential low quality cells from both the background drops and the cells as you would in any scRNAseq workflow (e.g. see Seurat tutorials and Luecken et. al. 2019 Mol Syst Biol).

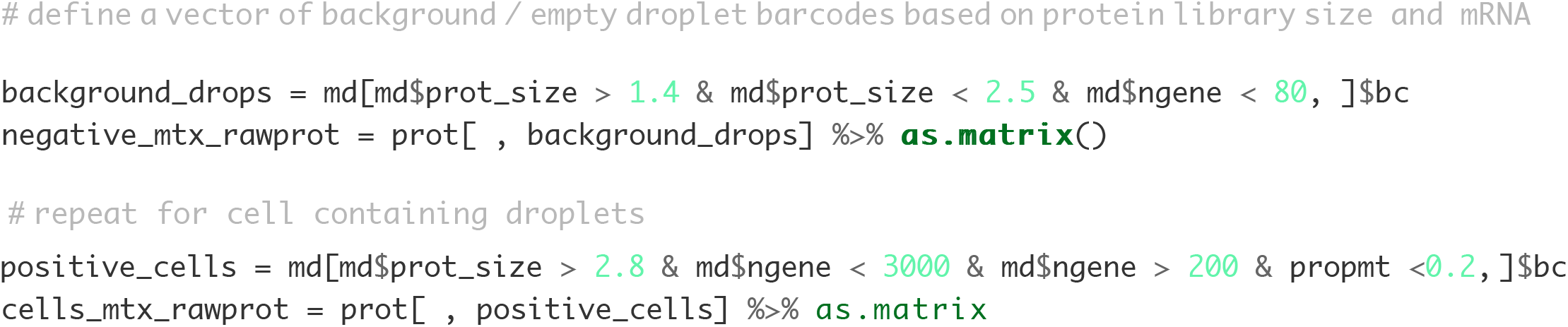

Check: are the number of cells in line with the expected recovery from the experiment?

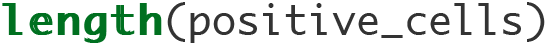

[1] 3481

Yes. After quality control above we have 3481 cells which is in line with the 5000 cells loaded in this experiment and closely matches the estimated cells from the filtered output from cell ranger (not shown). The protein library size distribution of the cells (orange) and background droplets (blue) used for normalization are highlighted below. >67,000 negative droplets are used to estimate the ambient background for normalizing the 3,481 single cells.

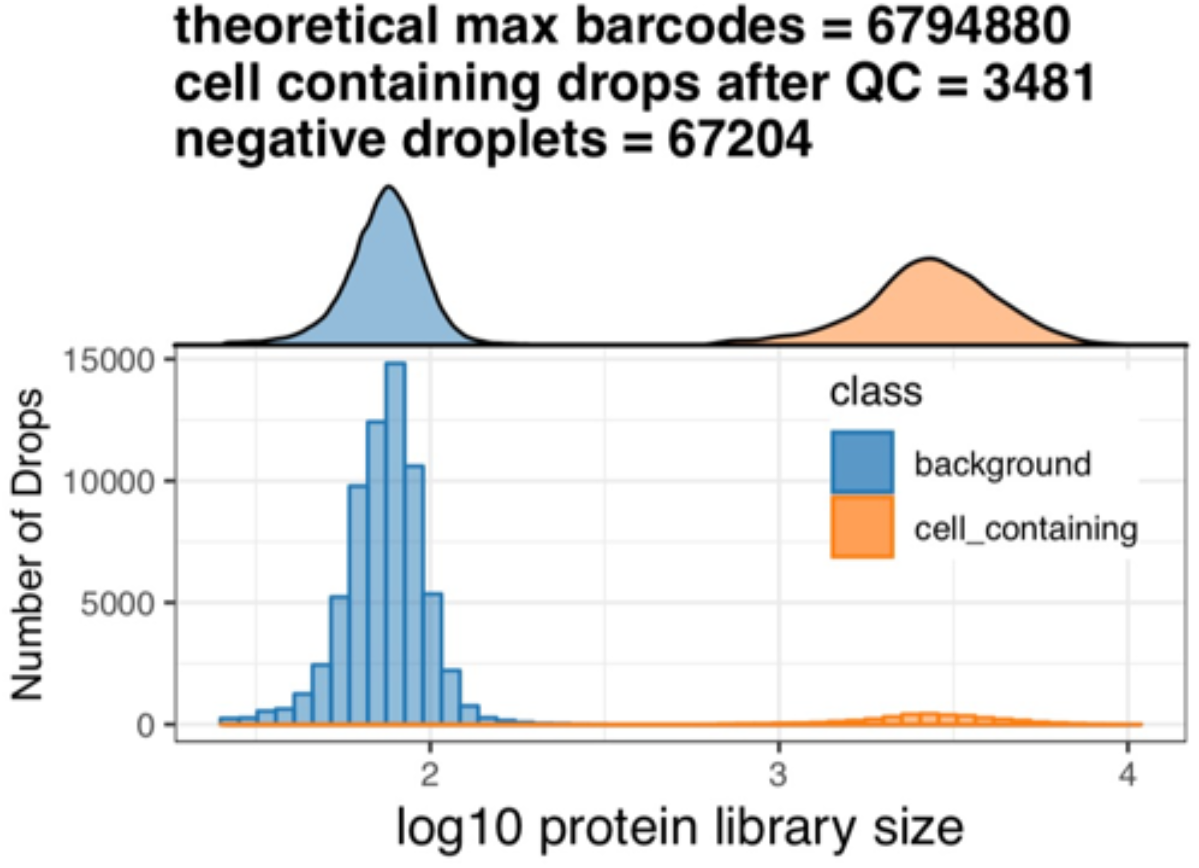

#### Step II: normalize protein data with the DSBNormalizeProtein Function

If you have isotype control proteins, set denoise.counts = TRUE and use.isotype.control = TRUE and provide a vector containing names of isotype control proteins (the rownames of the protein matrix that are isotype controls). For example, in this experiment, rows 30:32 of the data are the isotype control proteins so we set isotype.control.name.vec = rownames(cells_mtx_rawprot)[30:32]. If you don’t have Isotype controls see the section ‘Quick overview for experiments without isotype controls’.

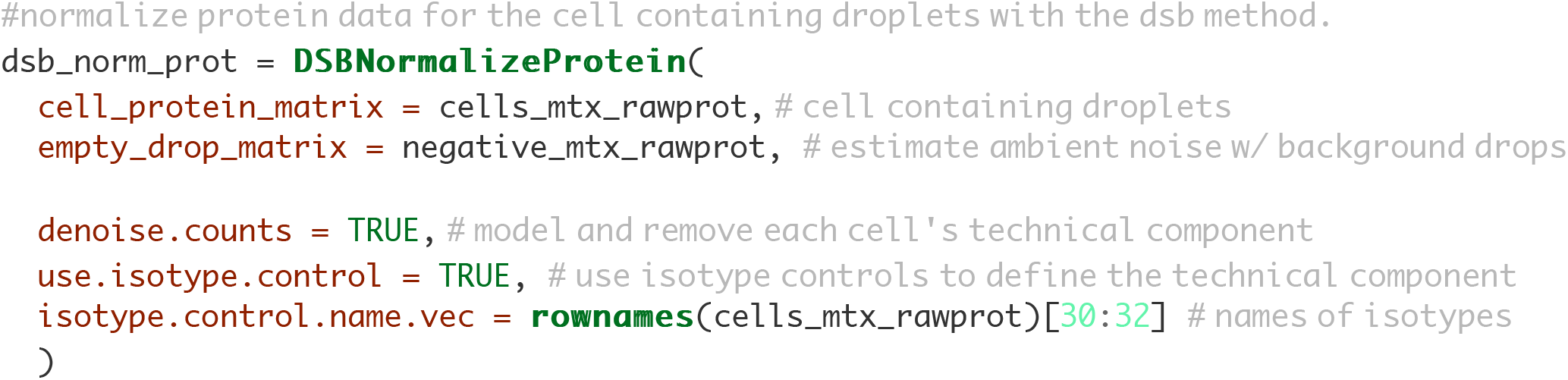

The function returns a matrix of normalized protein values which can be integrated with any single cell analysis software. We provide an example with Seurat, Bioconductor and Scanpy below.

### Integrated workflow with Seurat

#### Step III (option a) Integrate with Seurat

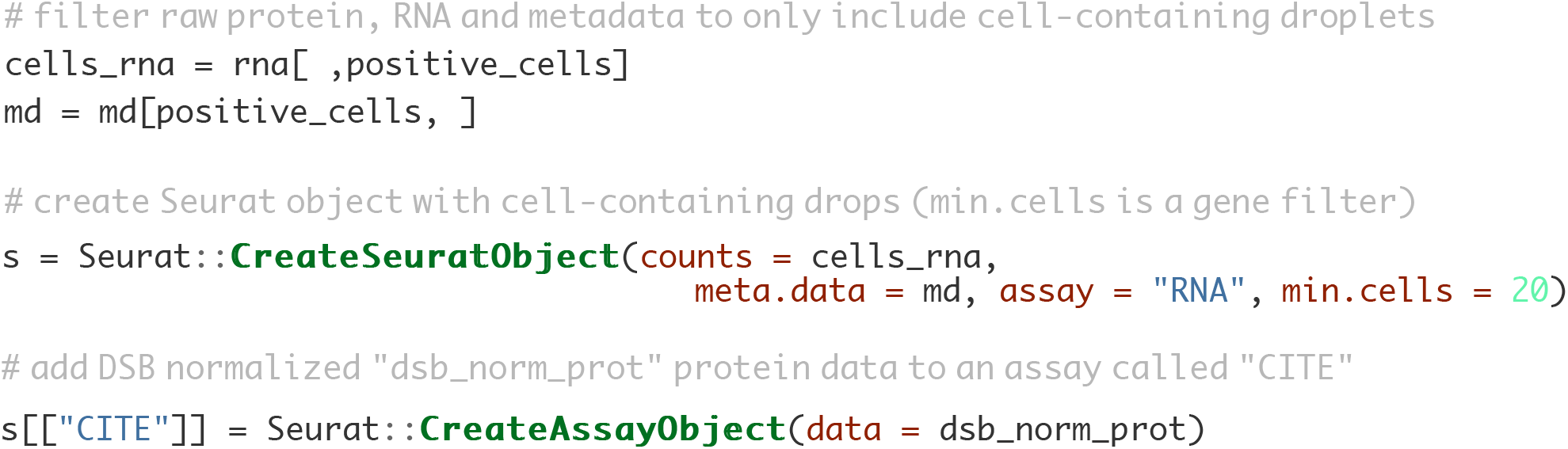

This object can be used in downstream analysis using Seurat.

#### Suggested Step IV: Protein based clustering + cluster annotation

This is similar to the workflow used in our paper Kotliarov et. al. 2020 Nat. Medicine where we analyzed mRNA states *within* interpretable clusters defined by dsb normalized protein data. We first run spectral clustering using S eurat directly on the dsb normalized protein values **without** reducing dimensionality of the cells x protein matrix with PCA.

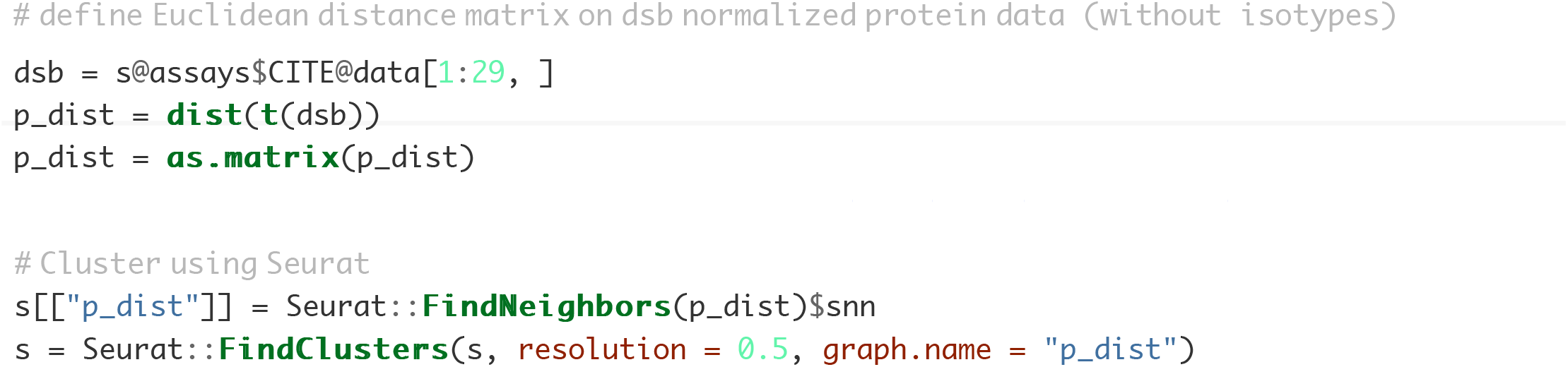

#### Suggested Step V: cluster annotation based on average dsb normalized protein values

A heatmap of average dsb normalized values in each cluster help in annotating clusters results. The values for each cell represent the number of standard deviations of each protein from the expected noise from reflected by the protein ‘s distribution in empty droplets, +/- the residual of the fitted model to the cell-intrinsic technical component.

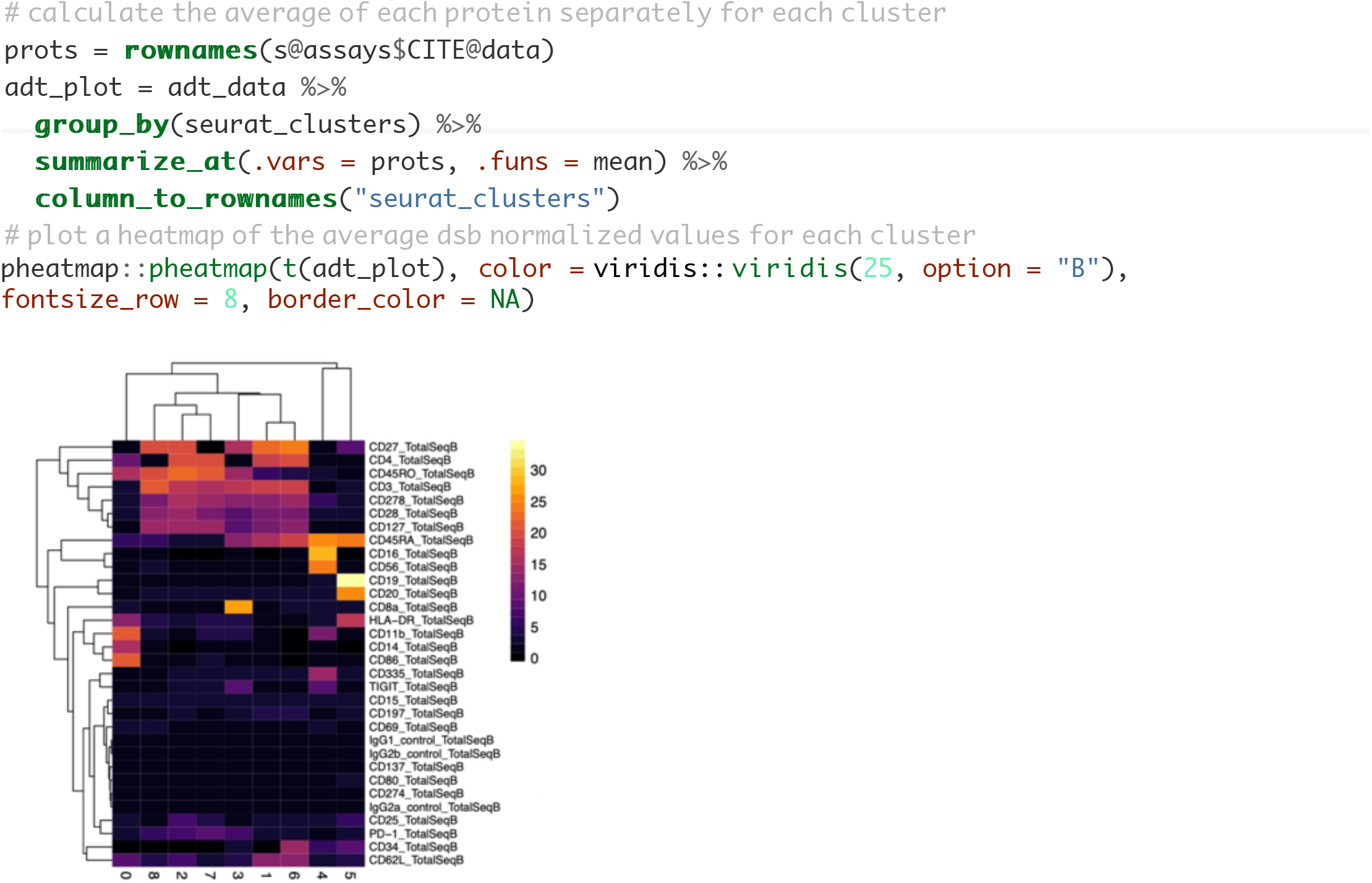

#### Visualization of single cell protein levels on the interpretable dsb scale

Dimensionality reduction plots can sometimes be useful visualization tools, for example to guide cluster annotation similar to the averaged values above. Below we calculate UMAP embeddings for each cell directly on the dsb normalized protein values-we recommend this rather than using principal components for tSNE or UMAP We then create a combined dataframe with the UMAP embeddings, dsb normalized protein values, and all cell metadata variables. This ‘tidy’ dataframe can be used for further statistical modeling or for visualization with base R functions or ggplot.

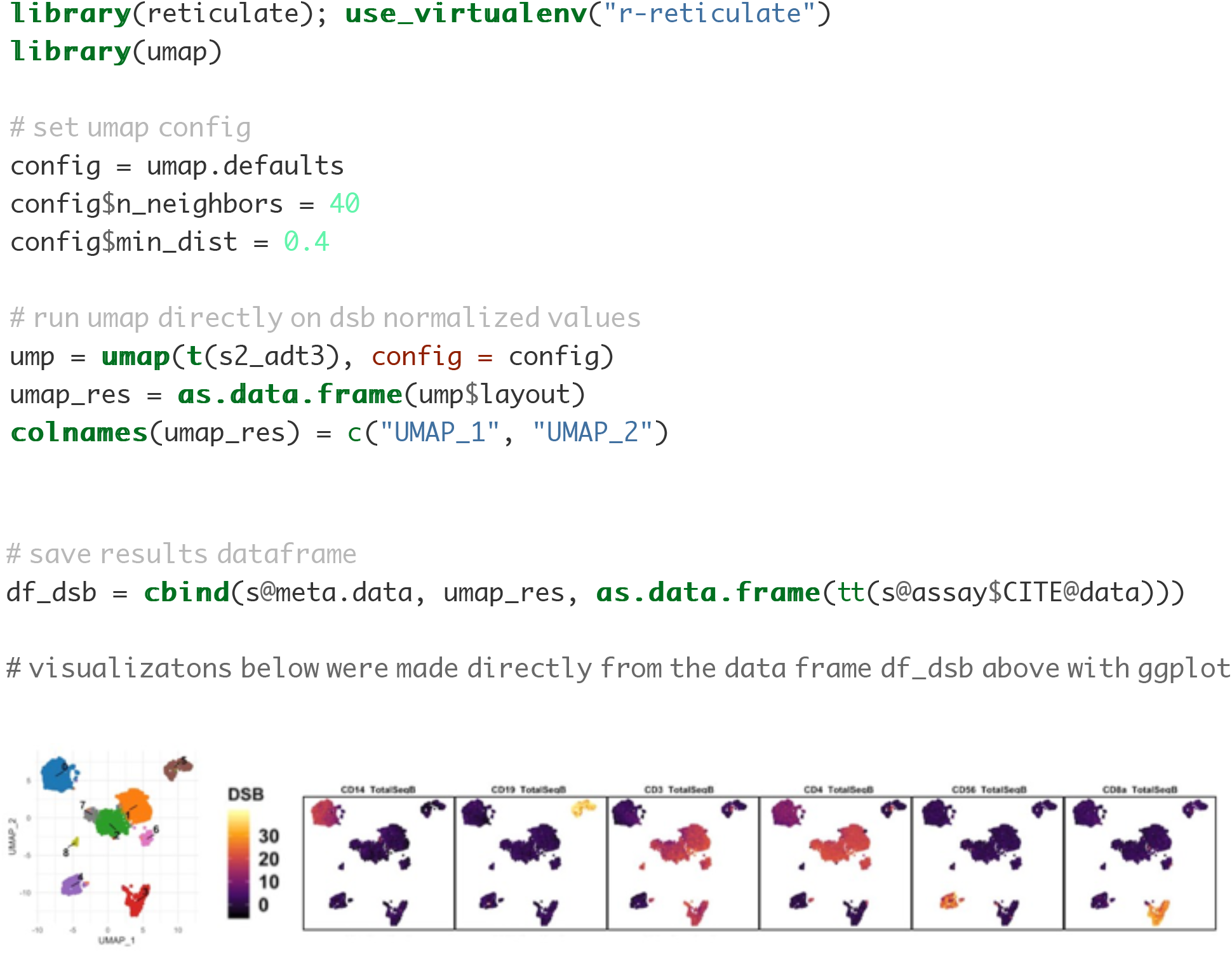

### Integration with Bioconductor and Scanpy

#### Step III (option b): integration with Bioconductor’s SingleCellExperiment class

Rather than Seurat you may wish to use the SingleCellExperiment class to use Bioconductor packages - this can be accomplished with this alternative to step III above using the following code. To use Bioconductor’s language, we store raw protein values in an ‘alternative Experiment’ in a

SingleCellExperiment object containing RNA counts. We add the dsb normalized protein matrix created in step II in the logcounts ‘assay’ of the protein ‘alternative experiment’.

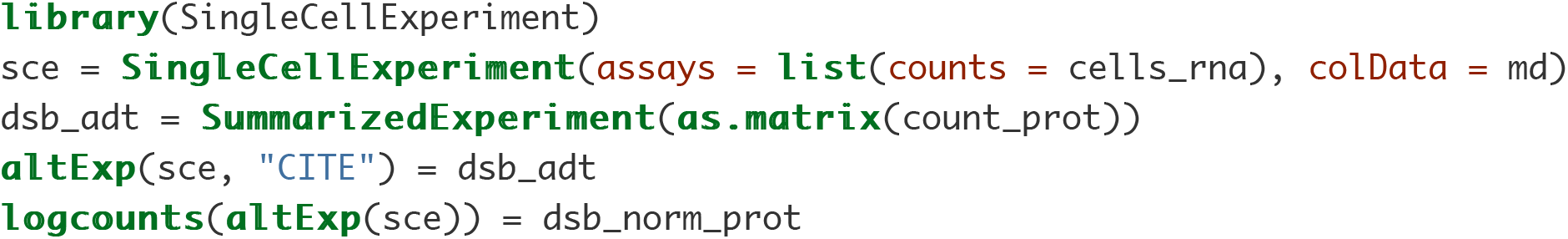

#### Step III (option c): integration with Scanpy using pythons AnnData class

You can also use dsb normalized values with the AnnData class in Python by following steps 1 and 2 above to load data, define drops and normalize with dsb in R. The simplest option is then to use reticulate to create the AnnData object from dsb denoised and normalized protein values as well as raw RNA data. That object can then be imported into a python session. Anndata are not structured as separate assays; we therefore need to merge the RNA and protein data. See the current Scanpy CITE-seq workflow and more on interoperability between Scanpy Bioconductor and Seurat

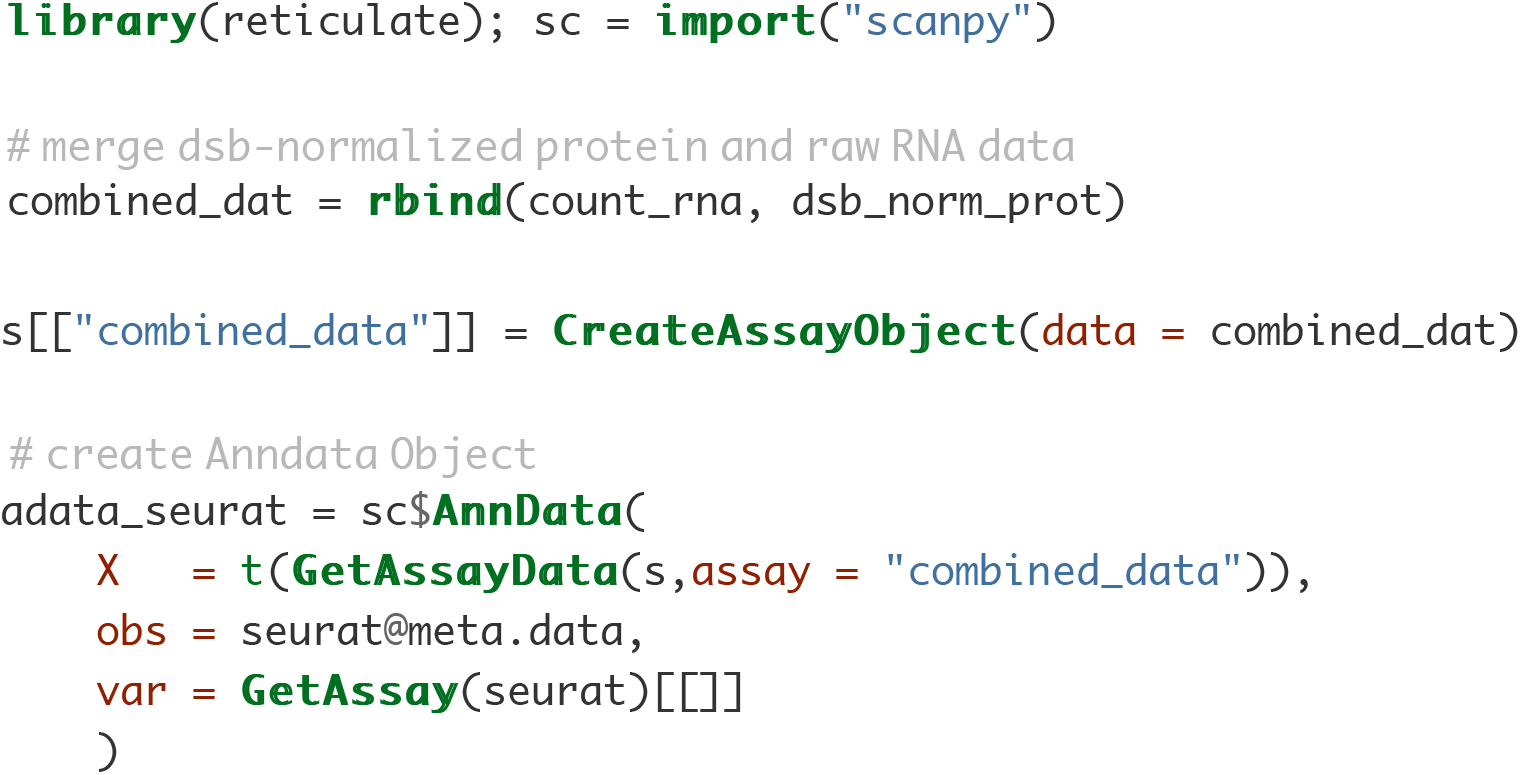

### Multiplexing experiments using barcoding antibodies or genomic barcoding

In multiplexing experiments with cell superloading, demultiplexing functions define a “negative” cell population which can then be used to define background droplets for dsb. In our data, dsb normalized values were nearly identically distributed when dsb was run with background defined by demultiplexing functions or protein library size (see the paper). HTODemux (Seurat) deMULTIplex (Multiseq) demuxlet

#### Example workflow using dsb with background defined by sample demultiplexing functions

As in step Ia we load the **raw** output from cell ranger. Prior to demultiplexing we use the min.genes argument in the Seurat::Readl0X function to partially threshold out some background drops yet still retain sufficient (often > 80,000 droplets per 10X lane depending on experiment) from which to estimate the background. This balances memory strain when demultiplexing tens of thousands of cells with requirements of the Seurat::HTODemux function to have sufficient empty drops to estimate the background population of each Hash antibody. Importantly, the HTODemux function may not converge if the cell ranger filtered output was loaded since there will not be sufficient negative drops to estimate the background for each hashing antibody. Increasing the number of drops used in demultiplexing will result in more droplets defined by the function as “negative” which can increase the confidence in the estimate of background used by dsb.

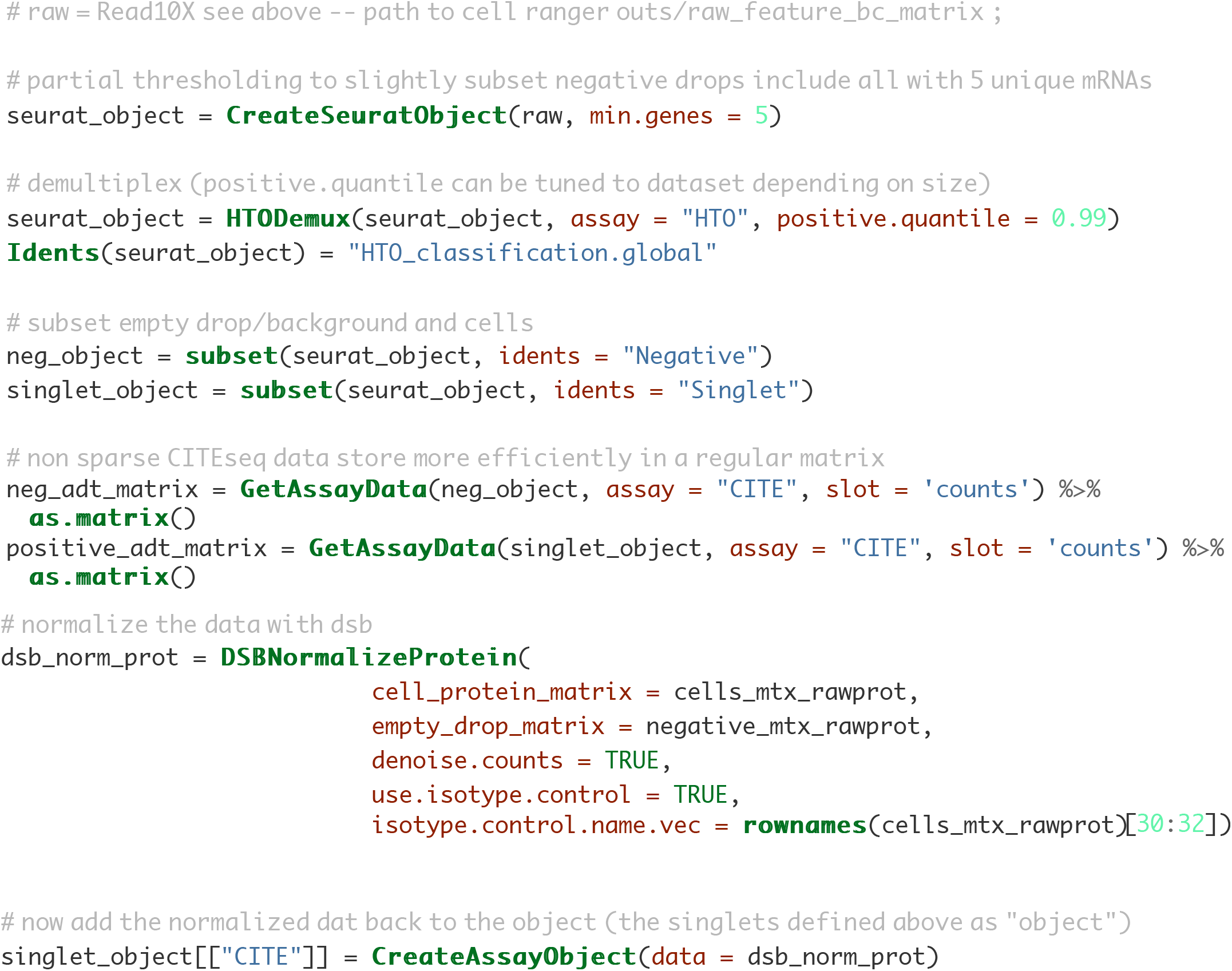

## Supplementary analysis code documentation

This documentatio n provides instructions for downloading all data used in th is analysis and for runnin g analysis scripts to reprod uce all results reported in the manuscript. The analysis code can be download ed at https://github.com/niaid/dsb_manuscript.

### Normalizing and denoising protein expression data from droplet-based single cell profiling

#### Analysis code to reproduce manuscript results

Mulè MP*, Martins AJ*, Tsang JS. **Normalizing and denoising protein expression data from droplet based single cell profiling**. bioRxiv. 2020;2020.02.24.963603.

This analysis pipeline reproduces all results and figures reported in the paper above. This paper describes control experiments and statistical modeling used to characterize sources of noise protein expression data from droplet based single cell experiments (CITE-seq, REAP-seq, Mission Bio Tapestri etc.) and introduces the dsb R package for normalizing and denoising droplet-based surface protein data.

The dsb method was developed in **John Tsang's Lab** by Matt Mulè, Andrew Martins and John Tsang.

The R package dsb is hosted on CRAN

link to latest dsb release on CRAN

link to dsb Github repository

**All data used below is available in analysis ready format at this figshare repository**: https://doi.org/10.35092/yhjc.13370915

#### Instructions for analysis workflow

This reproducible analysis workflow was run on a laptop with 16GB RAM. To run the analysis, 1) download the dsb_manuscript github repository, 2) download data from the figshare link above and add the /data folder directly to the root directory containing the .Rproj file; this directory should now contain dsb_manuscript.Rproj, the files readme.md and readme.rmd, the directory V2 and the directory you just added, data. One can view the commented code and run each script in each subdirectory as listed below, or source each R script in the order they appear below. No file paths need to be specified or changed. Each R script is self-contained, reading data from the /data folder and writing to figures or results files within each analysis subdirectory relative to the root directory through use of the the R package here.

#### software package versions used in this analysis

Please see full session info at the end of this script

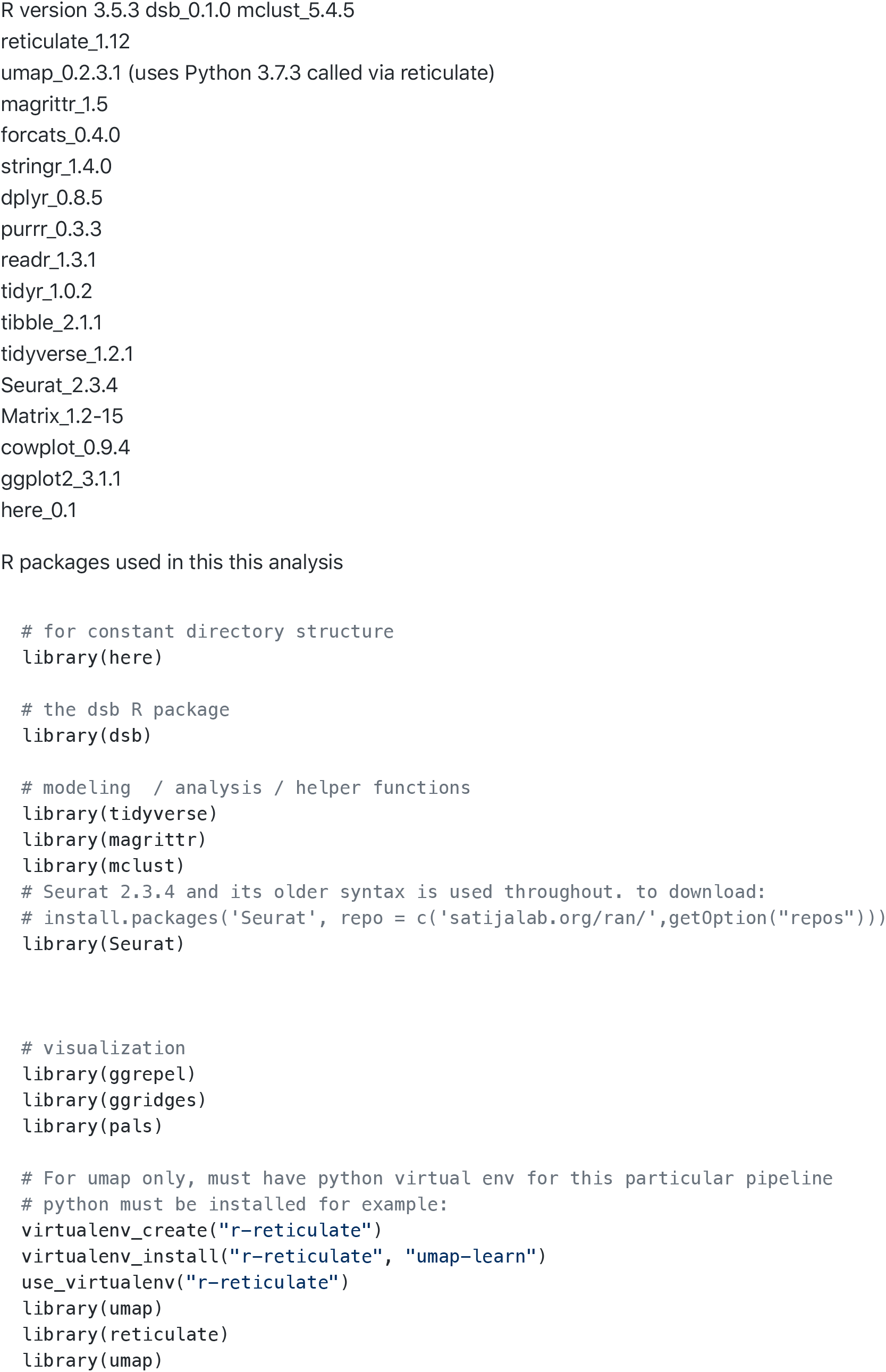

#### 3) Add downloaded data to the repository

After downloading the repository located at https://github.com/niaid/dsb_manuscript, add the data folder to the repository at the top level (where the file .rproj is located). Prior to running any analysis confirm the data are in the correct repository with the scripts below.

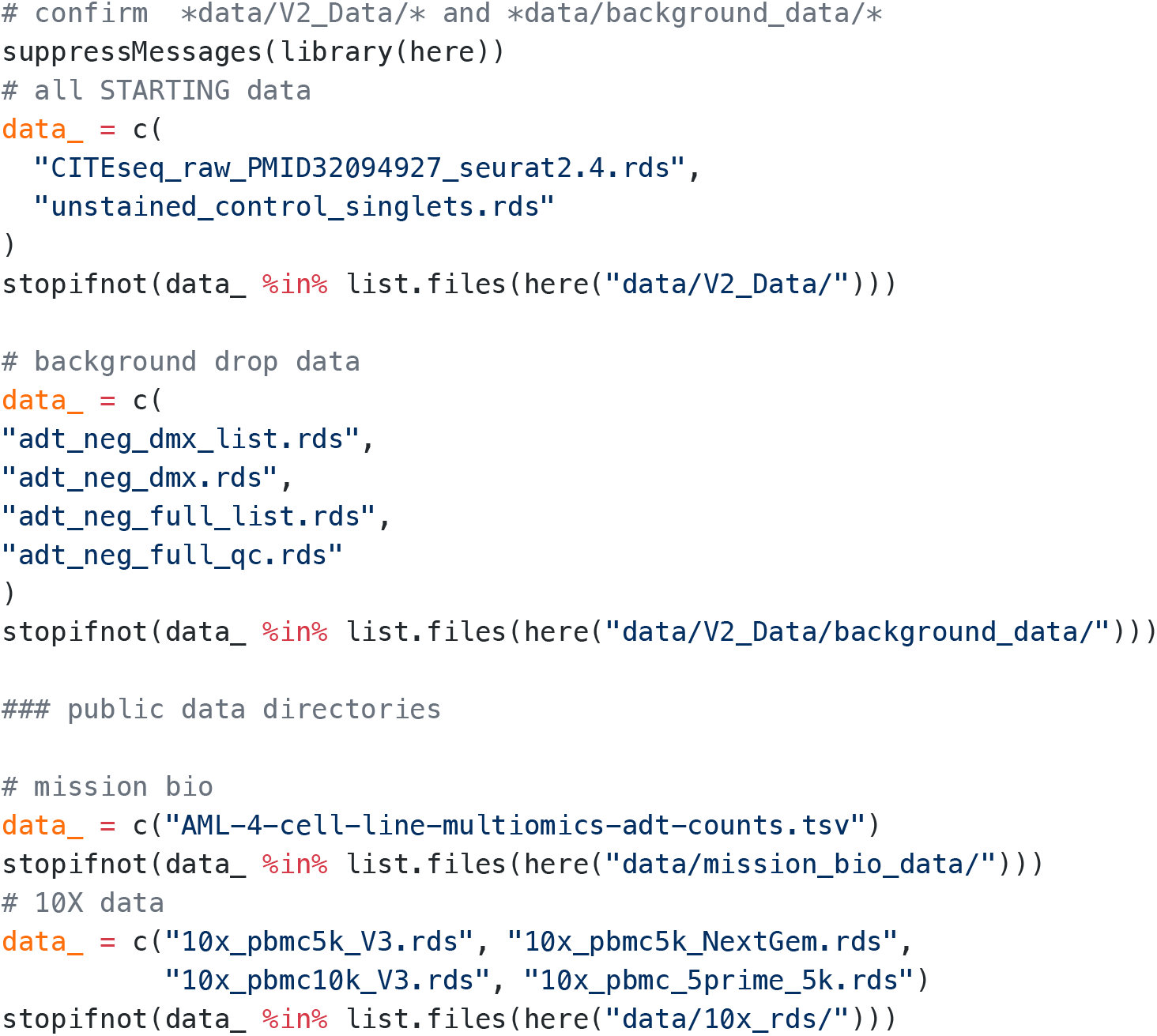

#### 4) Reproduce analysis results

These can be run line by line in an active R session or by sourcing the script. As described above, file paths do not need to be changed.

#### dsb normalize PBMC data from 20 individuals.

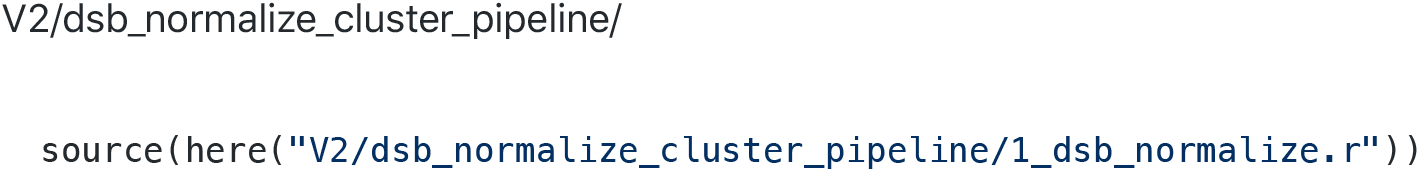

#### run umap

reads data from:

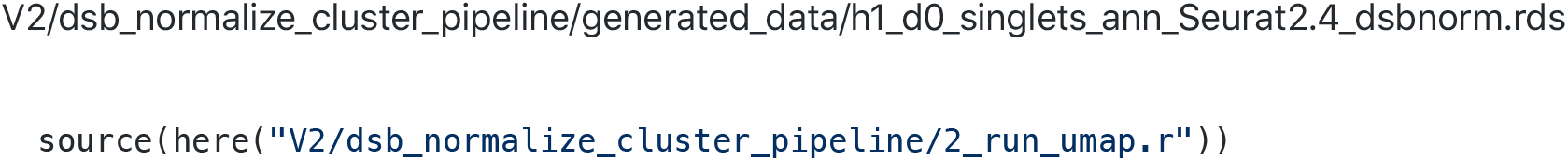

#### Figure generation and modeling results.

These scripts compare different normalizations with dsb and produce various comparison analysis and visualization between methods.

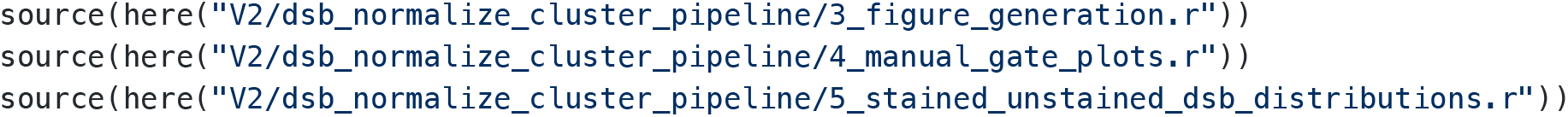

#### noise variable measurements and analysis of dsb underlying modeling assumptions

This section illustrates the dsb process in steps and models each step underlying the method. 6 and 6a analyze the correlation structure of variables comprising the technical component in step II of the dsb method. 7 models the correlation between multiple measurements of experimentally and model derived background noise with ‘ground truth’ background from unstained control cells spiked into the cell pool prior to droplet generation. In 8, the per-cell two component Gaussian mixture model is compared to k = 1-6 component models and the resulting single cell fits are analyzed.

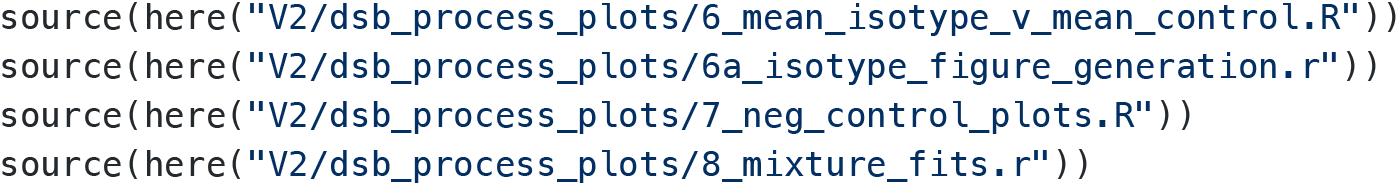

#### Multi vs single batch normalization and background drop sensitivity analysis

empty_drop_threshold_batch is an assessment of single vs multi batch normalization and sensitivity of each normalization scheme to defining background with hashing or library size distribution. mu1_noise_correlations is analysis of the robustness of μ1 background assessment and correlation with μ2 and isotype control means using 100 random sampling of μ1 proteins from each cell.

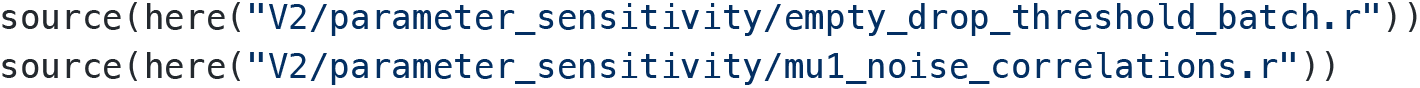

#### External 10X genomics data analysis: “NextGem”, “V3”, and “5 Prime” assays

These scripts are identical for each data set with tuned parameters at the beginning of each script. Read Cell Ranger raw output, select negative drops, run DSB normalization, run each normalization modeling step separately, cluster cells and plot distributions across clusters and on biaxial gates. Test underlying modeling assumptions for each external dataset.

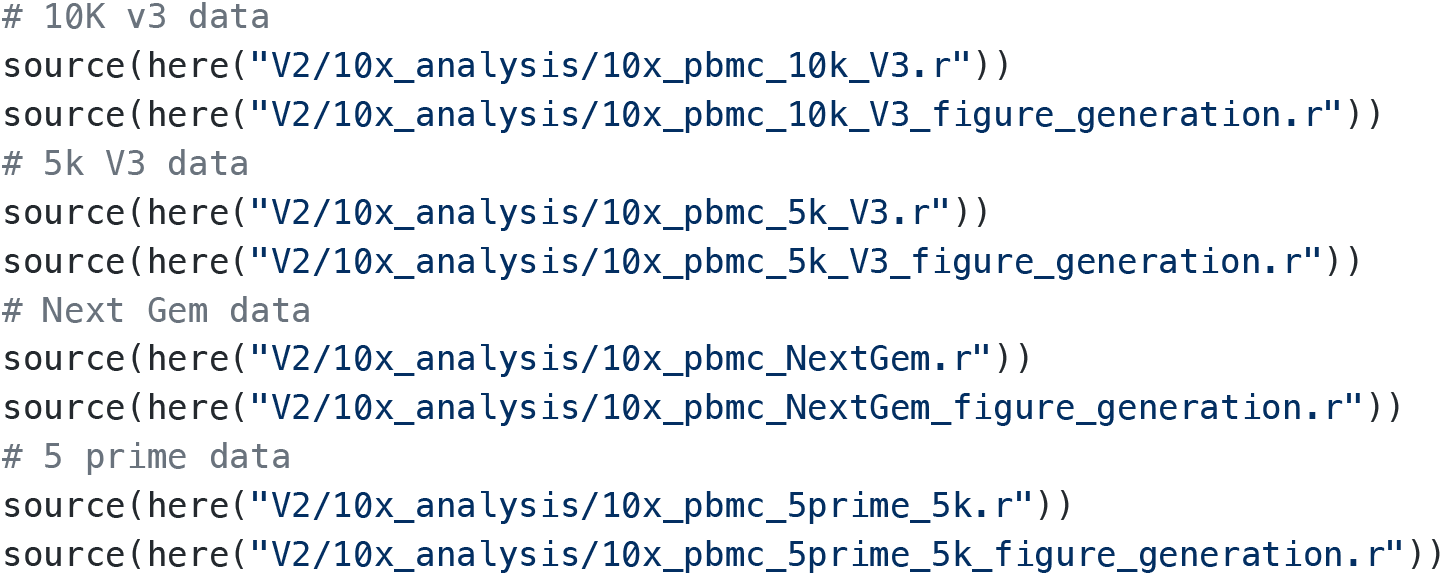

#### dsb normalize protein data from Mission Bio tapestri platform

The data downloaded from MissionBio are reformatted for dsb and normalized using dsb step I - ambient correction using empty droplets.

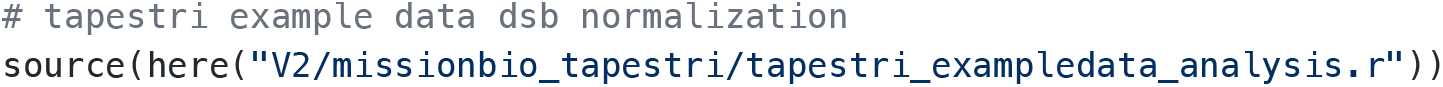

#### public dataset sources

Data are available in analysis-ready format at the figshare link above for convenience. For 10X data, downloaded only the the *raw, not the filtered data* in the links below downloaded **feature / cell matrix (raw)** output. dsb uses the non cell containing empty droplets that are in the raw output to estimate background - see package documentation.

Next GEM: https://support.10xgenomics.com/single-cell-gene-expression/datasets/3.0.2/5k_pbmc_protein_v3_nextgem

V3 (5K): https://support.10xgenomics.com/single-cell-gene-expression/datasets/3.0.2/5k_pbmc_protein_v3

V3 (10K): https://support.10xgenomics.com/single-cell-gene-expression/datasets/3.0.0/pbmc_10k_protein_v3

5’ https://support.10xgenomics.com/single-cell-vdj/datasets/3.0.0/vdj_v1_hs_pbmc2_5gex_protein

Mission Bio Tapestri: https://portal.missionbio.com/datasets/4-cell-lines-AML-multiomics download the following text file ‘AML-4-cell-line-multiomics-adt-counts.tsv’

#### initialization script for public datasets

This code is here for reference. Conversion of public 10X genomics datasets to .rds files.

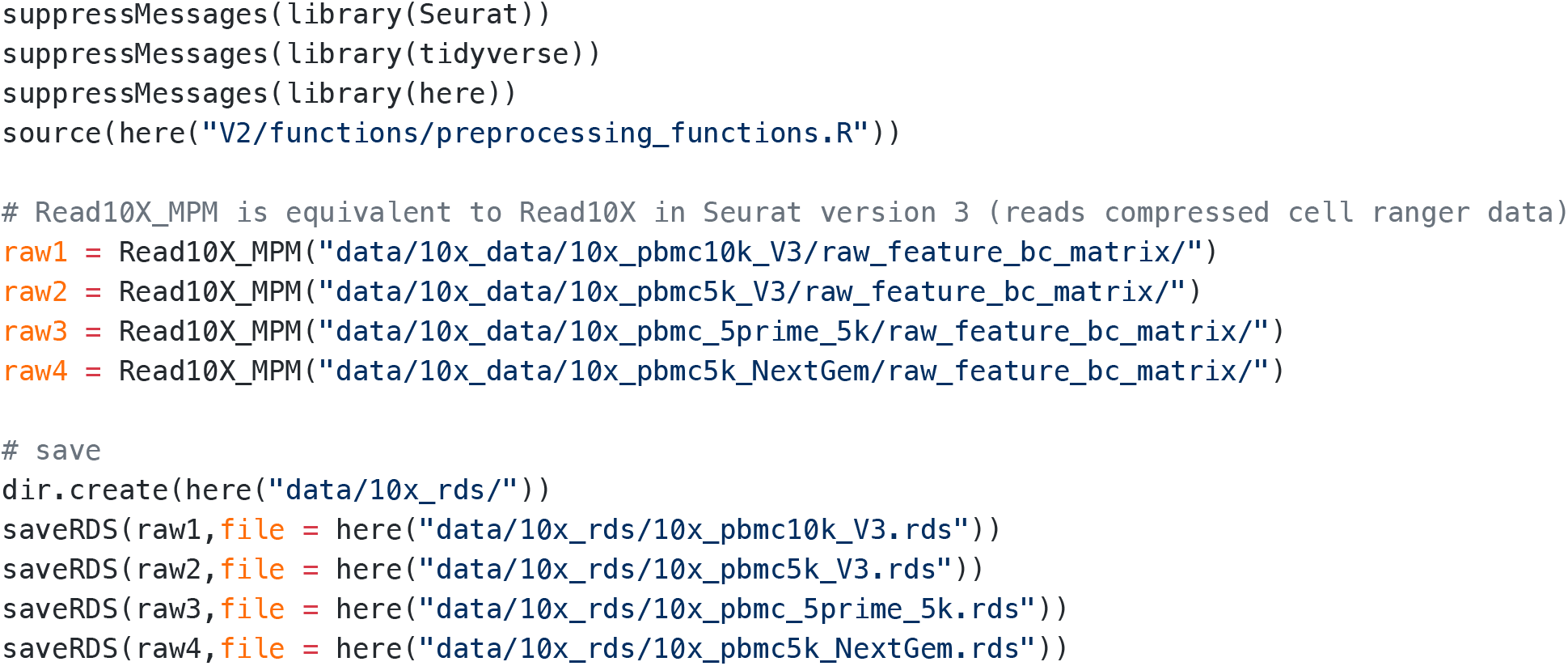

#### initialization script for healthy donor 53k cell CITE-seq data.

The code below was run outside of this workflow for convenience, it is shown here for reference. This step reformats the dataset hosted at the figshare repository that is associated with the manuscript. The script below removes the protein normalized data slot which used an earlier version of the DSB package for normalization. In addition, normalized RNA data, metadata, clustering snn graphs and tSNE results are removed to reduce object size and cell type annotations as reported in the paper linked above are added to the object as metadata.

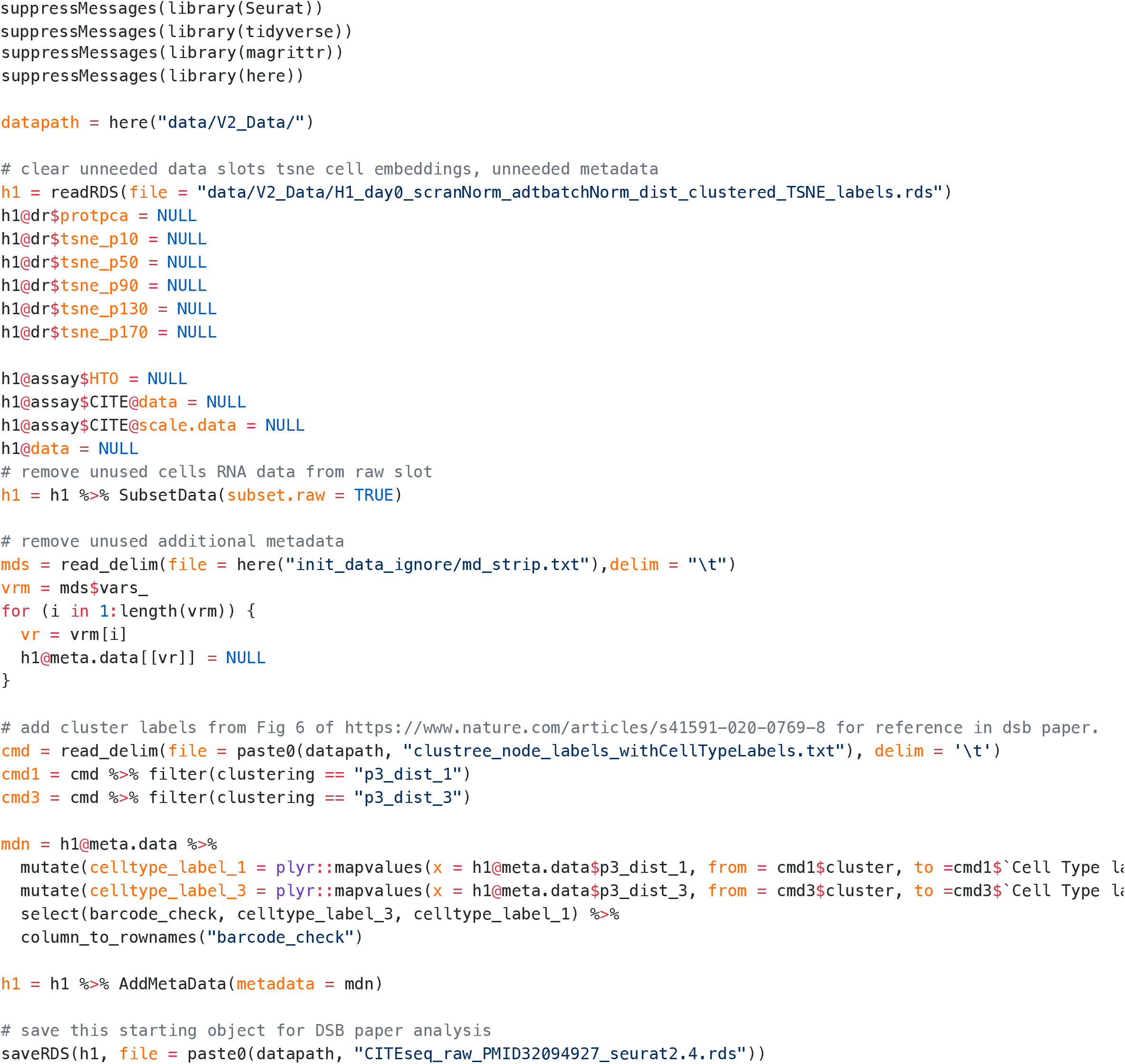

#### Project options and Sessioninfo

R version 3.5.3 attached base packages: stats graphics grDevices utils datasets methods base

other attached packages: mclust_5.4.5 reticulate_1.12 umap_0.2.3.1 magrittr_1.5 forcats_0.4.0 stringr_1.4.0 dplyr_0.8.5 purrr_0.3.3 readr_1.3.1 tidyr_1.0.2 tibble_2.1.1 tidyverse_1.2.1 Seurat_2.3.4 Matrix_1.2-15 cowplot_0.9.4 ggplot2_3.1.1 here_0.1

loaded via a namespace (and not attached): readxl_1.3.1 snow_0.4-3 backports_1.1.4 Hmisc_4.2-0 plyr_1.8.4 igraph_1.2.4.1 lazyeval_0.2.2 splines_3.5.3 inline_0.3.15 digest_0.6.25 foreach_1.4.4 htmltools_0.3.6 lars_1.2 rsconnect_0.8.16 gdata_2.18.0 checkmate_1.9.3 cluster_2.0.7-1 mixtools_1.1.0 ROCR_1.0-7 modelr_0.1.4 matrixStats_0.54.0 R.utils_2.8.0 askpass_1.1 prettyunits_1.0.2 colorspace_1.4-1 rvest_0.3.4 haven_2.1.0 xfun_0.7 callr_3.2.0 crayon_1.3.4 jsonlite_1.6 survival_2.43-3 zoo_1.8-6 iterators_1.0.10 ape_5.3 glue_1.3.1 gtable_0.3.0 pkgbuild_1.0.3 kernlab_0.9-27 rstan_2.19.3 prabclus_2.3-1 DEoptimR_1.0-8 scales_1.0.0 mvtnorm_1.0-10 bibtex_0.4.2 Rcpp_1.0.1 metap_1.1 dtw_1.20-1 htmlTable_1.13.1 foreign_0.8-71 bit_1.1-14 proxy_0.4-23 SDMTools_1.1-221.1 Formula_1.2-3 stats4_3.5.3 tsne_0.1-3 StanHeaders_2.21.0-1 htmlwidgets_1.3 httr_1.4.0 gplots_3.0.1.1 RColorBrewer_1.1-2 fpc_2.2-1 acepack_1.4.1 modeltools_0.2-22 ica_1.0-2 pkgconfig_2.0.2 loo_2.3.1 R.methodsS3_1.7.1 flexmix_2.3-15 nnet_7.3-12 tidyselect_0.2.5 rlang_0.4.5 reshape2_1.4.3 cellranger_1.1.0 munsell_0.5.0 tools_3.5.3 cli_1.1.0 generics_0.0.2 broom_0.5.2 ggridges_0.5.1 evaluate_0.14 yaml_2.2.0 npsurv_0.4-0 processx_3.3.1 knitr_1.23 bit64_0.9-7 fitdistrplus_1.0-14 robustbase_0.93-5 caTools_1.17.1.2 RANN_2.6.1 packrat_0.5.0 pbapply_1.4-0 nlme_3.1-137 R.oo_1.22.0 xml2_1.2.0 hdf5r_1.2.0 compiler_3.5.3 rstudioapi_0.10 png_0.1-7 lsei_1.2-0 stringi_1.4.3 ps_1.3.0 lattice_0.20-38 vctrs_0.2.4 pillar_1.4.1 lifecycle_0.1.0 Rdpack_0.11-0 lmtest_0.9-37 data.table_1.12.2 bitops_1.0-6 irlba_2.3.3 gbRd_0.4-11 R6_2.4.0 latticeExtra_0.6-28 KernSmooth_2.23-15 gridExtra_2.3 codetools_0.2-16 MASS_7.3-51.1 gtools_3.8.1 assertthat_0.2.1 openssl_1.4 rprojroot_1.3-2 withr_2.1.2 diptest_0.75-7 parallel_3.5.3 doSNOW_1.0.16 hms_0.4.2 grid_3.5.3 rpart_4.1-13 class_7.3-15 rmarkdown_1.13 segmented_0.5-4.0 Rtsne_0.15 lubridate_1.7.4 base64enc_0.1-3

